# Eco-Evolutionary Feedbacks Between Predator’s Linkage Disequilibrium and Prey Densities Maintain Diversity

**DOI:** 10.1101/523720

**Authors:** Swati Patel, Reinhard Bürger

## Abstract

Diversity occurs at multiple scales. Within a single population, there is diversity in genotypes and phenotypes. At a larger scale, within ecological communities, there is diversity in species. There have been a number of studies to investigate how diversity at these two scales influence each other through what has been termed eco-evolutionary feedbacks. Here we study a three-species ecological module called apparent competition, in which the predator is evolving in a trait that determines its interaction with two prey species. Unlike previous studies on apparent competition, which employed evolutionary frameworks with greatly simplified genetics, we study a multilocus eco-evolutionary model in which the predator’s evolving trait is determined by two diallelic loci. We ask how eco-evolutionary feedbacks with these two loci affect the coexistence of the prey species and the maintenance of polymorphisms within the predator species. In doing so, we uncover a novel eco-evolutionary feedback between the prey species and the linkage disequilibrium of the two loci of the predator. Through a stability analysis, we demonstrate how these feedbacks affect polymorphisms at both loci and, among others, may generate stable cycling.

## 1. Introduction

In recent years, there has been a growing body of literature devoted to understanding the feedbacks between ecological and evolutionary processes. Such work has demonstrated that these feedbacks can have important reciprocal effects. For example, the evolution of traits affects classical ecological questions, such as when and how species coexist or when interspecific diversity is promoted, and the ecology of species interactions affects classical evolutionary questions, such as how intraspecific diversity is maintained. Up to now, eco-evolutionary feedbacks have predominantly been studied in models developed with a simplified genetic framework, for instance when a trait of a focal species is determined by a single genetic locus (e.g., Wilson and Turelli, 1986), or by a quantitative trait, for which only the mean responds to selection (e.g., Schreiber et al., 2011; Patel and Schreiber, 2015; Cortez and Patel, 2017), or by adaptive-dynamics approaches which assume asexual inheritance (e.g., Abrams and Kawecki, 1999; Abrams, 2006*a*,*b*; Rueffler et al., 2005; Vasseur and Fox, 2011).

While these and other studies demonstrate the reciprocal effect between ecological and evolutionary dynamics, and provide insight into the resulting consequences, they constrain the possible evolutionary responses by ignoring the complex genetic architecture underlying most quantitative traits (Falconer and Mackay, 1996; Charlesworth and Charlesworth, 2010). Population genetics studies demonstrate that these complexities can have important impacts on the evolution and fitness of the species by affecting the distribution of the focal trait in multiple ways (for reviews, see Bürger, 2000; Johnson and Barton, 2005). Thus, it is natural to suspect that these genetic processes within one focal species cascade to affecting the community it lives in. Hence, to uncover a full picture of the implications of eco-evolutionary feedbacks, we need to begin digging deeper into details of the genetics processes within a species and analyze their impacts on inter- and intra-specific diversity. In our work, we begin this effort by analyzing an eco-evolutionary apparent competition model, in which the predator is evolving in a trait that is governed by two diallelic, recombining loci and determines the interaction with the two prey.

Apparent competition (Holt, 1977) is a well-documented community ecological module consisting of two prey species that negatively affect each other through a shared predator (Figure 1; for a recent review see Holt and Bonsall (2017)). In particular, by nurturing the shared predator, each species suppresses the other. This phenomenon has been widely observed in nature amongst a variety of ecosystems, including marine (Menge, 1995; Schmitt, 1987), tropical (Morris et al., 2004), and agricultural (Settle and Wilson, 1990; Muller and Godfray, 1997). Like many important ecological modules, such as exploitative competition, predator-prey, or intraguild predation, apparent competition has been the subject of theoretical studies to examine the mechanisms and potential impacts of eco-evolutionary feedbacks, when one or more species are evolving in a trait affecting species interactions.

**Figure 1.**
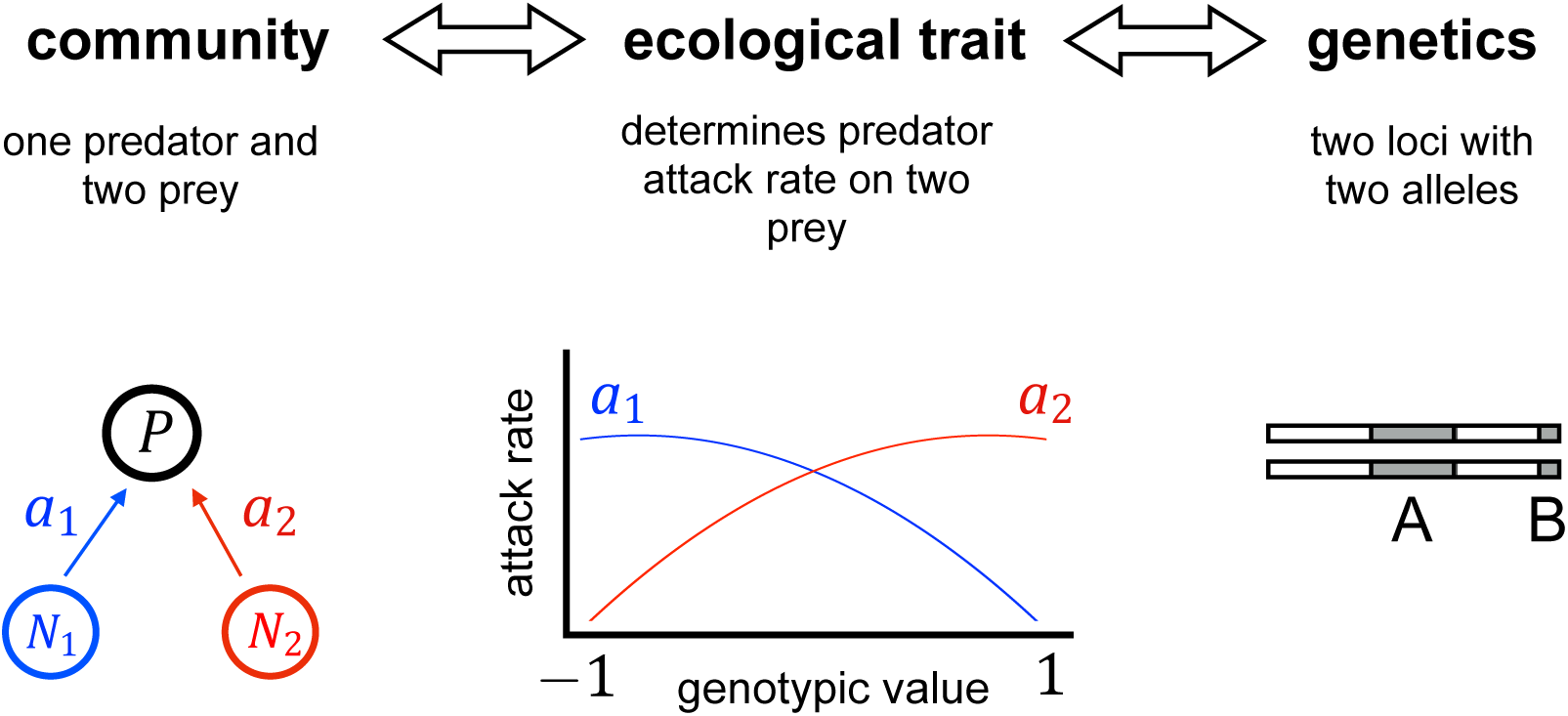
Schematic of the model. The three parts show the three-species community, how the attack rate of the predator on each prey depends on the genotype, and the two-locus genetics that determines the genotype value, i.e., the predator’s trait.

Previous studies have developed and analyzed a model in which an evolving predator has a trait governed by a single locus with two alleles that determines its consumption of the prey (Wilson and Turelli, 1986) and, additionally, how efficiently it uses what it consumes (Schreiber et al., 2018). Both studies show that, generally, genetic polymorphisms are more likely to be maintained with eco-evolutionary feedbacks, and thus intraspecific diversity is promoted.

On the other extreme, and considering a slightly different scenario, Schreiber et al. (2011) analyzed a model in which the shared predator evolves in a quantitative trait according to Lande’s (Lande, 1976) classical Gaussian model in which only the mean phenotype responds to selection, but the genetic and phenotypic variance, as well as the higher moments of the genotype and the phenotype distribution, remain constant. They found that eco-evolutionary feedbacks enable coexistence of the three species, i.e., interspecific diversity. In each of these studies, researchers attribute the maintenance of diversity to the following mechanism: as one prey increases in abundance, traits that strengthen the predators ability to consume that prey are selected for and this releases the less abundant prey from predation, which in turn generates an opposing selection pressure.

These and other studies were useful for building an understanding of basic mechanisms of eco-evolutionary feedbacks and their impacts. However, their simplified genetics ignores potentially important features of ecologically relevant traits which may often be polygenic. What makes the full dynamics of polygenic traits so difficult to study analytically is the interaction between selection and recombination. Selection generates statistical associations between alleles at different loci, so called linkage disequilibria, and recombination breaks up these associations. The latter may have the advantage of increasing the genetic variance, and thus accelerating the response to selection, but it also imposes a load by breaking up haplotypes carrying favorable allele combinations, thus reducing fitness. Additionally, epistasis, i.e., non-additive interactions amongst locus effects on traits or fitness, and differences in locus effects are common (Phillips, 2008) and may sometimes elevate or generate intraspecific variation (e.g., Gimelfarb, 1989; Nagylaki, 1989; Hastings and Hom, 1990; Gavrilets and Hastings, 1993; Bürger, 2000; Hermisson et al., 2003; Turelli and Barton, 2006; Pontz et al., 2018).

By assuming a constant variance, the quantitative genetics approach of Schreiber et al. (2011) does not allow for the variation to change as a consequence of evolution. In reality, allele frequencies and linkage disequilibria (see below) change in response to selection and thus, so does the genetic variance. In fact, numerous population-genetic studies show that these changes can be complex and are difficult to predict analytically (reviewed in Bürger, 2000). Whereas single-locus models admit changes in the variance, they do not account for the complexities of polygenic traits; in particular, the trait can assume only three values, which imposes serious constraints on the evolutionary dynamics.

The goal of this work is to investigate an analytically tractable model which allows for changes in the mean and the variance (as well as higher moments), and which exhibits sufficiently many different genotypes to grasp some of the intricacies of genuinely polygenic models. We aim to uncover the mechanisms and impacts of eco-evolutionary feedbacks when ecologically important traits are governed by multiple loci, and we explore how the underlying genetic architecture influences these mechanisms.

In particular, we extend the previous eco-evolutionary work in apparent competition to account for more complex genetic processes by examining the roles of epistasis, differences in locus effects, recombination, and linkage disequilibrium. Due to the analytical complexity of multilocus models, we restrict our study to two loci with two alleles. This model is sufficiently complex to capture these components of genetic architecture, while simple enough to enable analytical results and a mechanistic understanding.

We begin by introducing our new eco-evolutionary model which couples Lotka-Volterra population equations with classical two-locus population genetics equations. With this model, we first ask under what conditions is there coexistence amongst the species (intraspecific diversity)? Second, when does the predator population maintain polymorphisms (intraspecific diversity)? In these analyses, we explore the eco-evolutionary dynamics and show that it exhibits surprising features.

## 2. Model

In our eco-evolutionary model, we consider the joint dynamics of the densities *N*_1_ = *N*_1_(*t*) and *N*_2_ = *N*_2_(*t*) of two prey species, the density *P* = *P* (*t*) of their common predator, and the frequencies of the diploid predators’ genotypes producing a quantitative (phenotypic) trait (Figure 1). In the absence of predation, prey *k* experiences logistic growth with intrinsic growth rate *ρ*_*k*_ and carrying capacity *K*_*k*_. The predator population growth rate depends on its consumption of prey and each individual’s quantitative trait determines the attack rates on the two prey species. Usually, we suppress the dependence of the variables on time *t*.

The trait value of an individual predator is determined by two recombining diallelic loci. The effects of the alleles for locus one, labeled *A*_1_ and *A*_2_, are 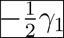 and 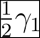, respectively, and for locus two, labeled *B*_1_ and *B*_2_, are 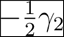 and 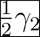, respectively. We label the four possible gametes *A*_1_*B*_1_, *A*_1_*B*_2_, *A*_2_*B*_1_, *A*_2_*B*_2_ by *i* = 1, 2, 3, 4, respectively, and denote the frequency at time *t* of gamete *i* by *p_i_*(*t*). The genotype *ij* consists of the gametes *i* and *j*, and the loci contribute additively to the genotypic value *g_ij_*, i.e., without dominance and epistasis. (Although we assume absence of epistasis among trait effects, there is still epistasis among fitness effects.) Thus, for instance, an individual with genotype *A*_1_*B*_1_*/A*_1_*B*_2_ has genotypic value 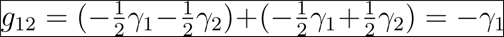. Here, *γ*_1_ and *γ*_2_ are both positive and, without loss of generality, we assume *γ*_1_ + *γ*_2_ = 1, and so each individual predator has a genotypic value between −1 and 1 (Figure 1). Because we ignore environmental contributions to the trait, the trait values equal the genotypic values *g*_*ij*_ and are given by

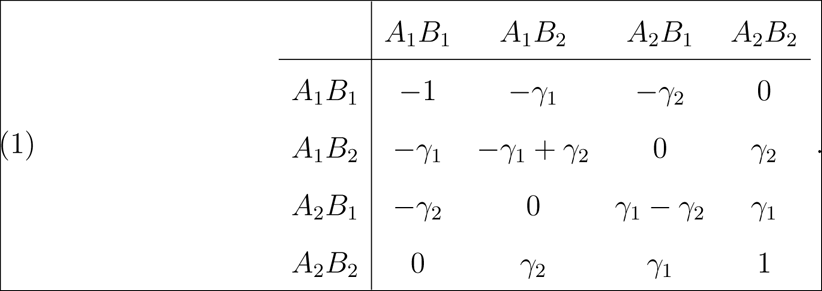

The attack rate of an individual with genotypic value *g* on prey *k* is a quadratic function given by

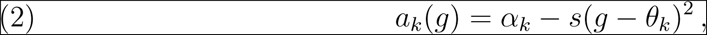

where we assume *θ*_1_ ≤ −1, *θ*_2_ *≥* 1, and *α*_*k*_ > *s*((−1)^*k*^ + *θ*_*k*_)^2^ for *k* = 1 and *k* = 2, so attack rates are positive and the optimal genotype for attacking prey 1 or 2 is − 1 or 1, respectively (Figure 1). The parameters *θ*_*k*_, *α*_*k*_, and *s* determine the shape of the attack rates and the induced trade-offs in efficiency of attacking one or the other prey. In particular, for fixed *θ*_*k*_ and *α*_*k*_, smaller *s* indicates a smaller range of possible attack rates and decreases the trade-off (Figure 2 solid curves). If *θ*_1_ = *−θ*_2_ is increased while *s* is decreased such that the range of attack rates stays the same, then smaller *s* decreases the concavity of the trade-offs (Figure 2 dotted curves). Hence, there is epistasis in the attack rates. We use these parameters to tune the shape of the trade-off and study the effects on the eco-evolutionary feedbacks.

Following Schreiber et al. (2011), we assume the fitness of an individual predator depends linearly on the attack rates and densities of the two prey. Hence, the fitness of a predator with genotype *g* is

**Figure 2.**
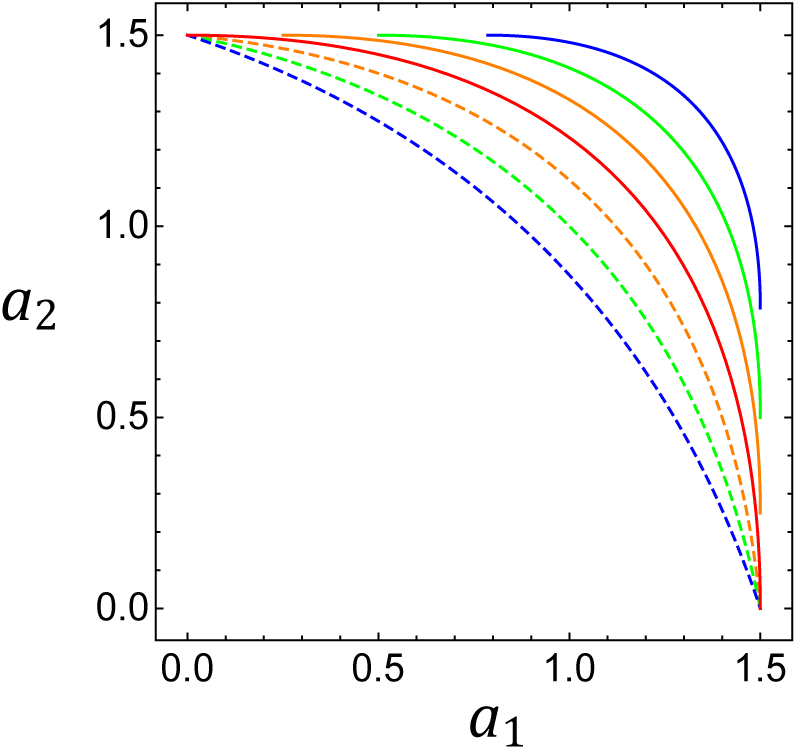
Trade-offs between attack rates on each prey. Each curve is a parametric plot of attack rate on prey 1 (*a*_1_) versus attack rate on prey 2 (*a*_2_) over the genotypic range from -1 to 1. Parameters are *α*_1_ = *α*_2_ = 1.5, and *s* = 0.375, 0.3125, 0.25, 0.1785, for red, orange, green, and blue, respectively. For solid curves, − *θ*_1_ = *θ*_2_ = 1, while for dotted curves *θ*_1_ = *θ*_2_ = 1, 1.2, 1.5 and 2.1, for red, orange, green, and blue, respectively, so that the range of attack rates stays constant and only the shape of the curve changes.

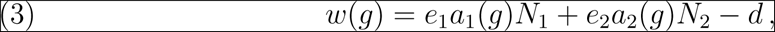

where *d* is the intrinsic death rate, and *e*_*k*_ is the conversion efficiency of prey *k*. Note that since the fitness of each genotype depends on the density of the two prey so does the strength of selection.

We model the evolutionary dynamics of the predator species by following the standard population genetics approach (e.g., Bürger, 2000). We assume that the predator species is diploid, mates randomly with respect to the trait, and the trait exhibits no sex differences. We also ignore random drift and mutation. Therefore, zygotes are in Hardy-Weinberg proportions and it is sufficient to follow gamete frequencies. We write *w*_*ij*_ for the fitness of genotype *ij*, where *w*_*ij*_ = *w*(*g*_*ij*_) is computed from (3) with *g*_*ij*_ as described above.

Approximating the standard population genetics discrete-time dynamics by a continuous-time dynamics, as is appropriate if both selection and recombination are weak (e.g., Nagylaki and Crow, 1974), the evolution of gamete frequencies is given by

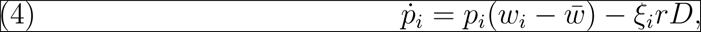

where *w*_*i*_ = Σ_*j*_*w*_*ij*_*p*_*j*_ is the marginal fitness of gamete *i*, 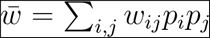 is the mean fitness of the predator population, *ξ*_1_ = *ξ*_4_ = *−ξ*_2_ = *−ξ*_3_ = 1, *r* is the recombination rate, and *D* = *p*_1_*p*_4_ *− p*_2_*p*_3_ denotes linkage disequilibrium.

For our purpose, it will be more convenient to describe the evolutionary dynamics by the alleles frequencies *p* = *p*_1_ + *p*_2_ of *A*_1_, *q* = *p*_1_ + *p*_3_ of *B*_1_, and *D* instead of the gamete frequencies. The reverse transformation is

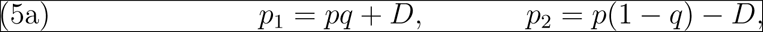

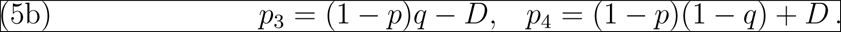

(This follows from the definition of *D* by utilizing that *p*_*i1*_ = *p*_*i*_ Σ_*j*_*p*_*j*_.) As is well known, the constraints *p*_*i*_ ≥ 0 and 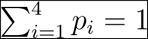 for *i* = 1,2,3,4,transform into

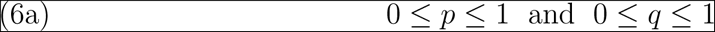

and

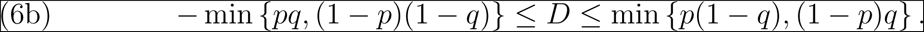

Altogether, this yields the coupled eco-evolutionary dynamics

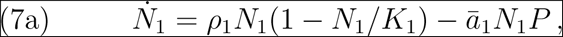

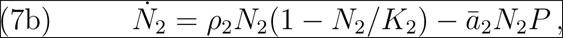

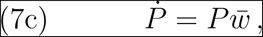

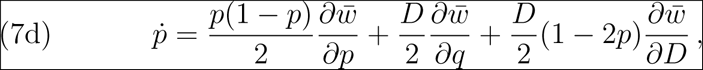

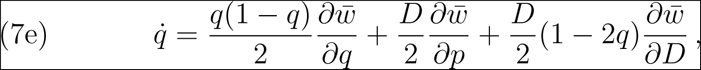

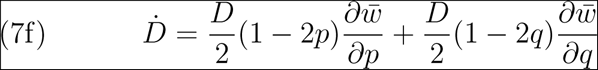

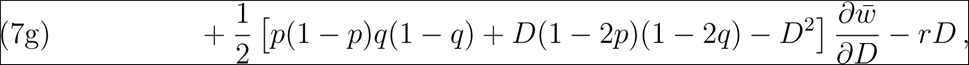

with

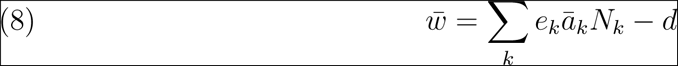

is the predator population mean fitness and

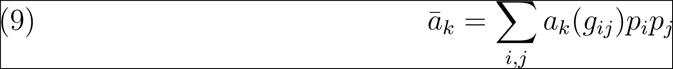

is the mean attack rate on prey *k*.

Full expressions with partial derivatives evaluated are given in Appendix 1. The first term in the expression for 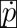 describes direct selection on locus A, which is frequency- and density-dependent because 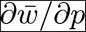 depends on population sizes as well as on *p* and *q* (see Appendix 1). The second term arises from indirect selection on locus *A* caused by the association (measured by *D*) with locus B. For 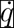, an analogous description applies. For 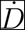, the first two terms describe the effects of the additive component of selection, the first term in the second line is due to epistatic effects (otherwise, 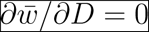), and the last term quantifies the erosion of linkage disequilibrium by recombination.

Straightforward computations show that mean genotypic value of the trait is

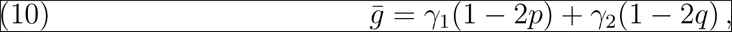

and the genetic variance is (e.g. Bürger 2000, p. 49)

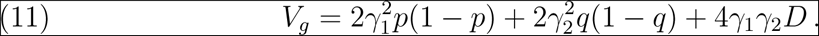

This shows that for given allele frequencies *p* and *q*, positive linkage disequilibrium inflates the variance, and negative linkage disequilibrium deflates it. Using 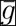 and *V*_*g*_, we can express the mean attack rate on prey *k* as a function of the allele frequencies

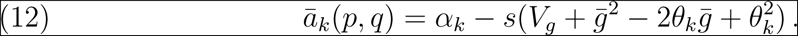

With this model, to understand how eco-evolutionary feedbacks affect interspecific diversity, we first derive conditions to ensure the coexistence of the two prey along with the shared predator (Section 3). Then, we ask how eco-evolutionary feedbacks affect the maintenance of polymorphisms. To do so, we partition our analysis into two parts: in part one (Section 4.1), we assume that recombination is strong and in part two (Section 4.2), we assume recombination is weak.

## 3. Results: Species Coexistence

We begin by analyzing when the two prey species coexist with one another despite the shared predator. Throughout all of our analyses, we assume that the carrying capacities of each prey are large enough to support the persistence of the predator 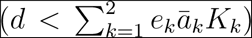 for all values of 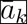 possible given the genotypes. We make this assumption since, when there is no predator to evolve, the community dynamics are uninteresting because each prey approaches its own carrying capacity. We find that the two prey coexist if

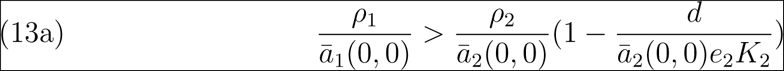

and

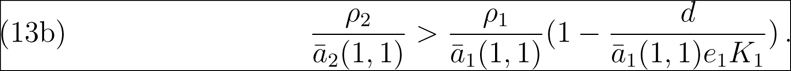

For derivations of these conditions, see Appendix 2. Heuristically, when prey 1 is rare and prey 2 is abundant, alleles *A*_2_ and *B*_2_ will be selected for in the predator, driving the entire population towards extreme genotypes close to 1, i.e., *p* and *q* to zero. Hence, the first condition ensures that prey 1 has positive growth rate when rare. Analogously, the second condition ensures that prey 2 has positive growth rate when rare. Together these give that the two species coexist provided both prey are initially present.

When the carrying capacities are large, the conditions (13) can be approximated as

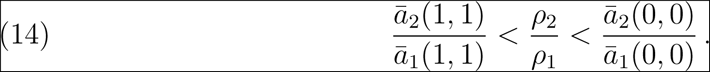

Hence, the two prey will coexist if the ratio of the intrinsic growth rate of prey 2 to 1 is between the ratios of the predator’s respective attack rates at the extreme genotypes. Condition (14) is always satisfied when growth rates are equal and attack rates are symmetric with respect to the predator genotypic values (as in Figure 1). Furthermore, when *θ*_1_ = −1 and *θ*_2_ = 1, then (14) simplifies to which illustrates the dependence on the selection parameter *s* (Figure 3). Increasing *s* strengthens the trade-off between attacking one prey versus the other and allows for coexistence (solid lines in Figure 2). Notice also that coexistence does not depend on recombination. In the following sections, we assume that condition (13) is met and ask what the effects are on the maintenance of intraspecific diversity, i.e., polymorphisms within the predator.

**Figure 3.**
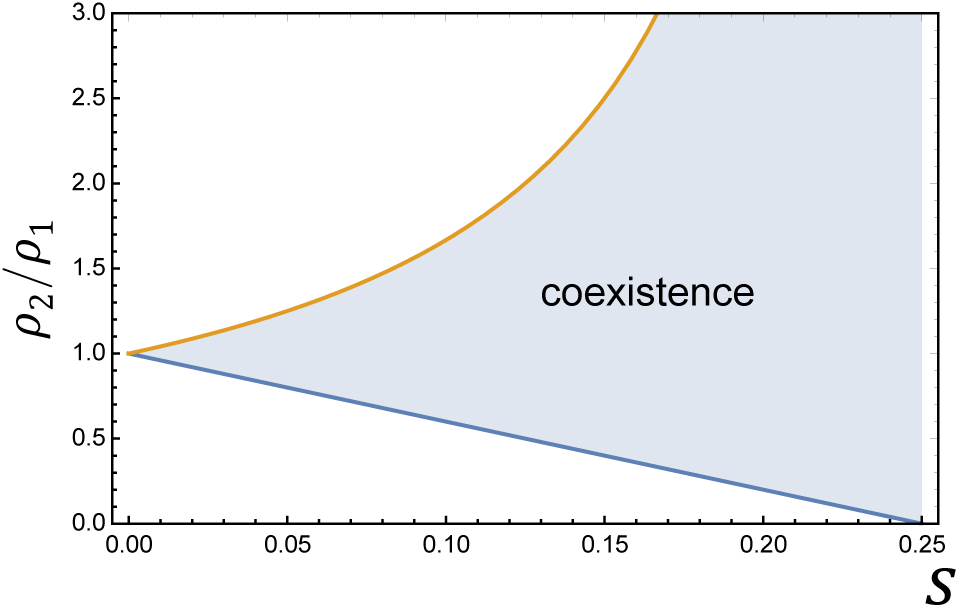
Regions of coexistence of two prey. The shaded region shows the values of *ρ*_2_*/ρ*_1_ and *s* for which symmetric prey species coexist, given a sufficiently large carrying capacity. The upper and lower bound of this region are given by the right and left term, respectively, in (15). Parameters are *α*_1_ = *α*_2_ = 1.

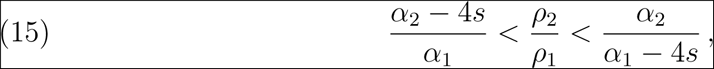

## 4. Results: Maintenance of polymorphisms

To investigate how eco-evolutionary feedbacks affect the maintenance of the polymorphism, we consider two assumptions: in Section 4.1 the loci are independent, and in Section 4.2 they are linked with some *r ≥* 0. Note that the former case arises as a limit of the latter when *r → ∞* (recall that *r* is the recombination rate, not the recombination probability). In each case, we begin by assuming both prey are symmetric.

By this, we mean that the two prey are equivalent in the absence of the predator and the attack rates of the predator on the two prey are symmetric with respect to the predator genotypes, i.e.,

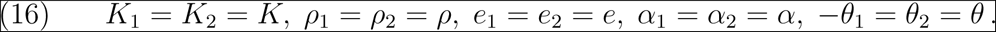

As a consequence, the attack rates satisfy *a*_1_(−*g*) = *a*_2_(*g*) (Figure 1). We make this assumption to avoid prey differences so we can focus on the effects of the predator’s genetic dynamics. This assumption also enables the derivation of clear analytical results. Then, we drop this symmetry assumption and study how the results change when the prey have different growth rates.

With the assumption of symmetric prey, we find that there is at least one equilibrium point at which all three species coexist and at which both loci are polymorphic. Such an equilibrium is obtained by setting 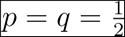, *N*_1_ = *N*_2_, and solving the equilibrium conditions for *N*_1_, *P*, and *D* under the assumptions *N*_1_ *>* 0, *P >* 0 and 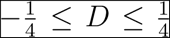. The (uniquely determined) expressions for the equilibrium values 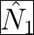 and 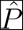 are complicated and given in Supporting Information 1 and in the *Mathematica* notebook in Supporting Information 2, and that for the linkage disequilibrium is given below in eq. (18b). We call this the fully polymorphic, symmetric equilibrium.

To understand the maintenance of diversity, we first analyze the stability of this fully polymorphic, symmetric equilibrium and then investigate the eco-evolutionary dynamics numerically using a numerical solver in *Mathematica*. The stability analysis provides insight into the eco-evolutionary feedbacks that affect the maintenance of intraspecific diversity.

Local stability of equilibrium points is determined by the stability modulus of the associated Jacobian matrix, *s*(*J*), which is the largest of the real parts of the eigenvalues. When *s*(*J*) *<* 0, the equilibrium is asymptotically stable, and when *s*(*J*) *>* 0, it is unstable. Typically, evaluating stability of six-dimensional systems is analytically cumbersome, and only really feasible through computational methods. However, we identified a way to greatly simplify the problem by a change of variables. In particular, we define

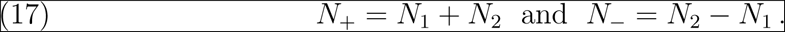

Therefore, instead of looking at the two prey densities individually, we will examine the total prey density *N*_+_ and the difference *N*_−_ between the two prey densities. With this change of variables and the ordering (*N*_+_*, P, D, p, q, N*_−_), the symmetric equilibrium becomes

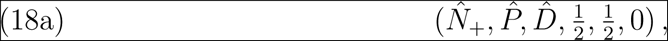

where

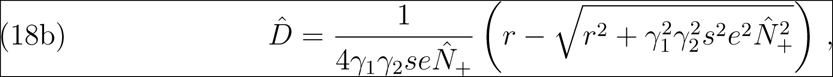

and 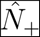 is given in Supporting Information 1, eq. (19.5). The Jacobian at this equilibrium now takes the simpler form of a block diagonal matrix:

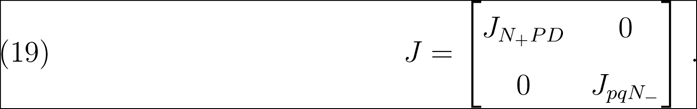

Here, *J*_*N*+*PD*_ and *J*_*pqN*−_ are a submatrix of the total prey density, predator density, and linkage disequilibrium and a submatrix of the two allele frequencies and the difference between the two prey densities, respectively. The exact expressions of the submatrices are given in Supporting Information 1 and the derivation in the *Mathematica* notebook in Supporting Information 2.

This change of variables is enlightening for two reasons. First, the block diagonal structure of the Jacobian spotlights two sets of critical eco-evolutionary first-order feedbacks (that is, how components affect other components) that affect stability: (1) a feedback between allele frequencies at both loci and the difference between the two prey densities, and, intriguingly, (2) the total prey density, predator density, and linkage disequilibrium (Figure 4). This suggests that, locally around the equilibrium point, the eco-evolutionary dynamics can be partitioned into these two uncoupled groups and examined separately.

**Figure 4.**
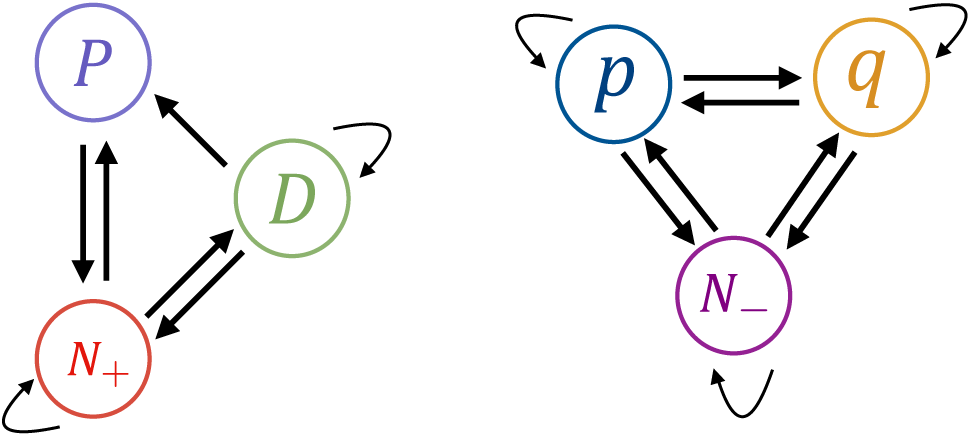
Feedbacks affecting stability of the fully polymorphic, symmetric equilibrium. The relevant first-order feedbacks inferred from non-zero entries of the Jacobian matrix at the fully polymorphic, symmetric equilibrium given in (18). Notice that the system is decoupled into two separate groups of variables.

Upon further examination of the submatrices, we observe higher-order feedbacks (that is, how components affect the *feedbacks* between other components). Interestingly, the predator density, total prey density, and the linkage disequilibrium influence the feedbacks between the allele frequencies and the prey density difference (Figure S1). In contrast, the feedbacks between the predator density, total prey density, and the linkage disequilibrium are unaffected by the allele frequencies and the prey density difference. In a subsequent section, we discuss how these higher-order feedbacks affect the dynamics.

Second, by reducing a six-dimensional Jacobian matrix into two, more tractable, three-dimensional submatrices (*J*_*N+PD*_ and *J*_*pqN*−_), we are able to derive analytical results for the stability conditions. Namely, we get two sets of conditions, one associated with each set of feedbacks (*s*(*J*_*N+PD*_) *<* 0 and *s*(*J*_*pqN−*_) *<* 0). Full expressions of the conditions are given in Supporting Information 1. For local stability, both conditions must be met. Additionally, when these two conditions are met, numerical solutions suggest that the equilibrium is globally stable. Conversely, if either condition is not satisfied, then the equilibrium is unstable and we can determine which of the two ecoevolutionary feedbacks is driving instability. Furthermore, in this case, the bifurcation patterns of *J*_*N+PD*_ and *J*_*pqN*−_ provide insight into the ensuing eco-evolutionary dynamics when the equilibrium is unstable.

### 4.1 Independent loci

When the two loci are independent, they are in linkage equilibrium (*D* = 0). By imposing *D* = 0, the eco-evolutionary dynamics of (7) is reduced to a 5-dimensional system because (7g) becomes void. In addition, (7d) and (7e) simplify substantially.

If additionally the prey are symmetric, as we assume in the sequel, then there is a fully polymorphic, symmetric equilibrium point at which all species coexist. It is given by 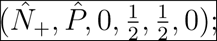; see Supporting Information 2 for full expressions. We find that, since 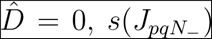 is always positive, and hence, this equilibrium is always unstable, due to the feedbacks between allele frequencies and the difference between the prey densities (Supporting Information 1). Furthermore, numerically solving for equilibrium points suggests that this is the unique fully polymorphic equilibrium with all three species coexisting.

Trivially, there are exactly four equilibria in which both loci are monomorphic, i.e., *p* = *q* = 0, *p* = 1 *− q* = 0, 1 *− p* = *q* = 0, and *p* = *q* = 1, and in which all three species might coexist (see Supporting Information 2 for full expressions). In addition to these equilibria, there may be other equilibria in which one locus is polymorphic, which we find numerically.

To infer the eco-evolutionary dynamics, we use the following visualization on the two-dimensional allele-frequency plane ((*p, q*); Figure 5). First, we use previous results that in the absence of evolutionary trait changes, the solutions of the ecological equations in (7) always approach a unique globally stable equilibrium (Takeuchi and Adachi 1983). For each pair of allele frequencies (and associated average attack rates), we determine the unique globally stable population equilibrium, and note whether it supports all three populations (coexistence) or just one prey and one predator, which are the only possibilities under our assumptions. This divides the (*p, q*) plane into three distinct regions, which indicate the community that is reached if allele frequencies remain within these bounds (shading in Figure 3 indicates coexistence).

**Figure 5.**
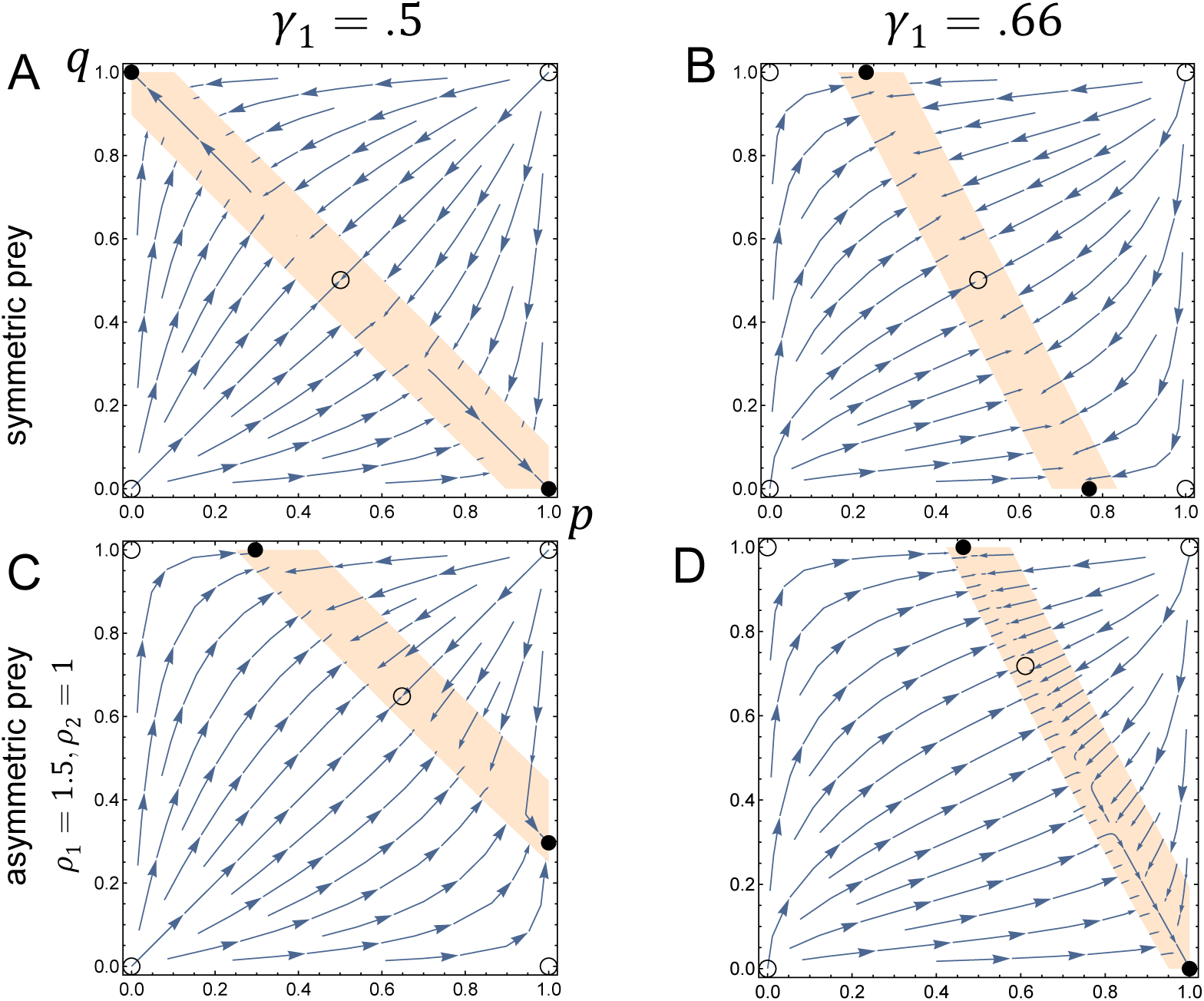
Eco-evolutionary vector fields when loci are independent. The shaded region delimits the range of allele frequencies for which all three species would coexist in the absence of any evolutionary change. This separates the region of allele frequencies for which prey 2 excludes prey 1 (upper right) and prey 1 excludes prey 2 (lower left). The arrows indicate directional vectors assuming population densities are at their corresponding equilibrium. Filled and open circles are stable and unstable, respectively, equilibrium points. Parameters in common for all panels are *α*_1_ = *α*_2_ = 0.5, −*θ*_1_ = *θ*_2_ = 1.1, *s* = 0.1, *e*_1_ = *e*_2_ = 1, *K*_1_ = *K*_2_ = 50, *d* = 2. For (A) *ρ*_1_ = *ρ*_2_ = 1, *γ*_1_ = *γ*_2_ = 0.5, (B) *ρ*_1_ = *ρ*_2_ = 1, *γ*_1_ = 0.66, *γ*_2_ = 0.34, (C) *ρ*_1_ = 1.5, *ρ*_2_ = 1, *γ*_1_ = *γ*_2_ = 0.5, and (D) *ρ*_1_ = 1.5, *ρ*_2_ = 1, *γ*_1_ = 0.66, *γ*_2_ = 0.34.

We then solve for the equilibrium population densities and determine the evolutionary dynamics (arrows in Figure 5) when the population densities are at equilibrium. From this, we predict the eco-evolutionary dynamics. While this is a projection of five dimensions onto two, hence may not always give a full picture of the dynamics, numerical solutions of the full dynamics agree with these predicted dynamics. Indeed, as suggested by the two-dimensional visualization, we observe numerically that there is a bistability with these two boundary equilibria. In other words, the predator evolves to one of the boundary equilibria. Importantly, this shows that when loci are independent, polymorphisms cannot be maintained at both loci.

Whether a polymorphism is maintained at one locus and, if so, which one, depends on the locus effects. When the two prey are symmetric and the loci have equal effects (i.e., *γ*_1_ = *γ*_2_ = 0.5), the two stable equilibria are such that both loci are monomorphic and alleles with opposing effects are fixed (Figure 5A). At both of these equilibria, both prey have equal density and the mean genotype is at its optimum of zero. When the two prey are symmetric and the loci contribute unequally to the genotype, the two stable equilibria are such that the locus with major effect is polymorphic and the locus with minor effect is monomorphic (Figure 5B).

Through numerical analysis, we examine the maintenance of polymorphisms when prey are asymmetric in intrinsic growth rate (but still within the bounds for prey coexistence given in (13)). We find that there is still one fully polymorphic equilibrium and that it is unstable. The predator evolves to the equilibrium point on the boundary with mean genotype better suited for consuming the prey with higher growth rate. When the two loci have equal effects, this promotes polymorphisms in one of the two loci (Figure 5C). Here, which locus maintains polymorphism depends on the initial allele frequencies as well as population densities. However, the number of polymorphic loci does not. If loci contribute unequally to the trait, then another scenario is possible: the predator either evolves to a monomorphic equilibrium or a single locus polymorphic equilibrium (Figure 5D).

### 4.2 Linked loci

When the two loci are not independent, then they are in linkage disequilibrium. In particular, beneficial combinations of alleles become more prevalent in the population. Recombination will break down this linkage and, hence, plays a critical role in the eco-evolutionary dynamics. If the prey are symmetric, i.e., (16) holds, then all three species coexist at the fully polymorphic, symmetric equilibrium (18), at which linkage disequilibrium is negative.

We analyze the stability of this fully polymorphic, symmetric equilibrium using both sets of feedbacks and determine the impacts of recombination and of locus effects. Additionally, we compare these stability conditions to the stability conditions in the absence of eco-evolutionary feedbacks to determine the role of feedbacks on the maintenance of diversity.

We find that when the locus effects are equal (i.e., *γ*_1_ = *γ*_2_ = 0.5) the fully polymorphic, symmetric equilibrium is always unstable (Figure 6; Supporting Information 1), in line with previous theory based only on the evolutionary dynamics, i.e., for stabilizing selection that is neither frequency- nor density-dependent (Wright, 1935; Bürger, 2000, Chap. 6.2). Furthermore, the equilibrium is unstable for unequal locus effects, i.e., *γ*_1_ ≠ *γ*_2_, when recombination rates are sufficiently large: by breaking up beneficial allele combinations, recombination creates a higher variance in fitness and thereby, imposes a destabilizing pressure on the double polymorphism (Figure 6; Supporting Information 1). However, if the locus effects are unequal and the recombination rate is below a threshold, the equilibrium is stable (Figure 6; Supporting Information 1). Hence, there is a critical recombination rate for stability and for maintenance of intraspecific diversity. Again, this finding is analogous to that for the frequency- and density-independent case (e.g., Gavrilets and Hastings, 1993; Bürger, 2000, Chap. 6.2).

**Figure 6.**
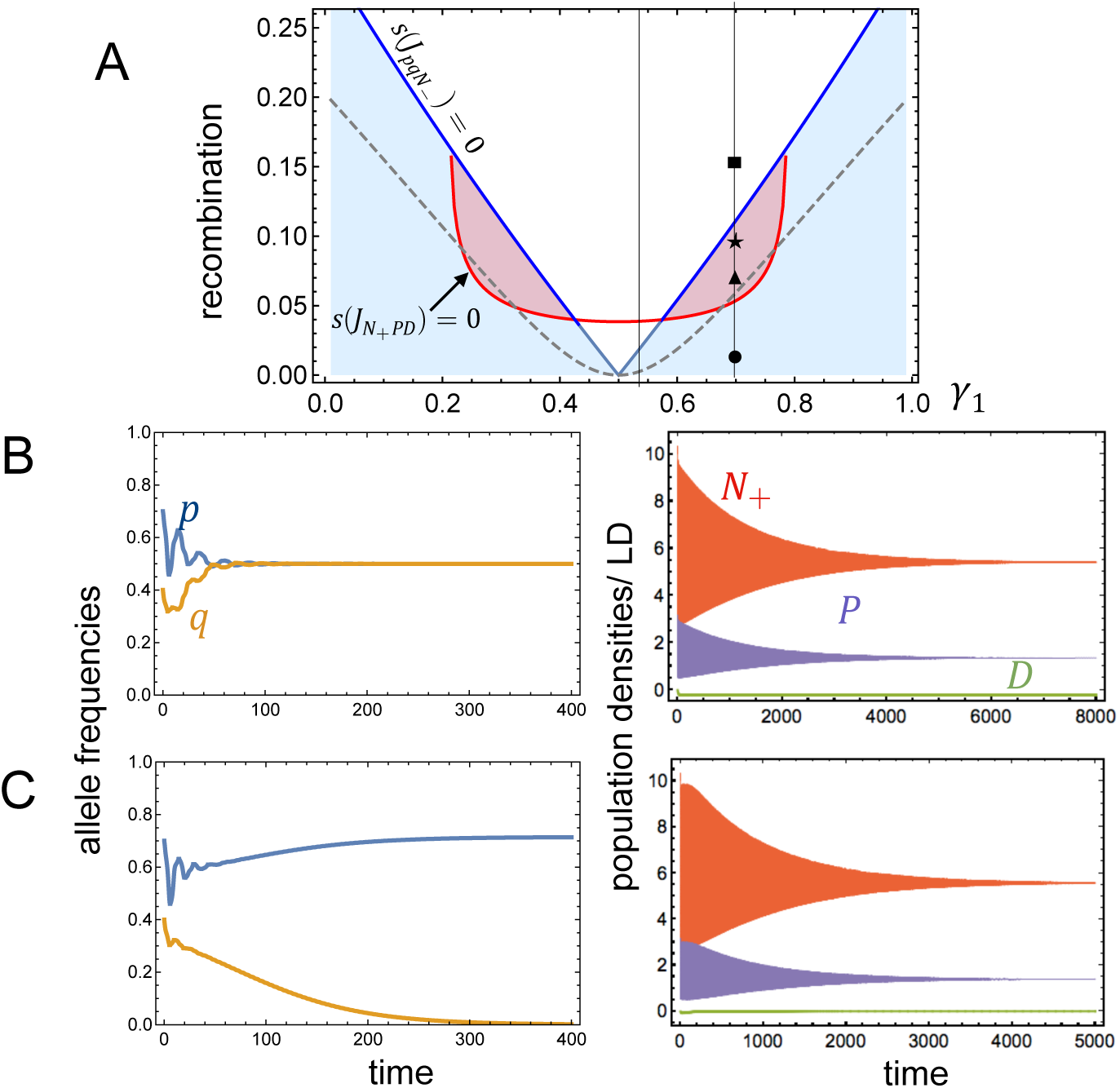
Stability of the fully polymorphic symmetric equilibrium. (A) Bifurcation diagram showing the dependence of the critical recombination rate for stability of the equilibrium (18) on the locus effects (*γ*_1_). The red curve shows *s*(*J_N_*_+*P D*_) = 0, and the blue curve shows *s*(*J_pqN−_*) = 0. The equilibrium is asymptotically stable below both curves (blue shading). Between the blue and the red curves (red shading), numerics show stable limit cycles (see Figures 7, 8). The dashed gray curve shows when the pure evolutionary dynamics (with no ecological feedback) changes stability. Parameters are *α*_1_ = *α*_2_ = 1, −*θ*_1_ = *θ*_2_ = 1.1, *s* = 0.2, *e*_1_ = *e*_2_ = 0.5, *K*_1_ = *K*_2_ = 1500, *ρ*_1_ = *ρ*_2_ = 1, *d* = 2. (B) Numerical solutions for our model with parameters as *γ*_1_ = 0.7 and *r* = 0.01, corresponding to the filled circle in (A). (C) Numerical solutions for our model with parameters *γ*_1_ = 0.7 and *r* = 0.15, corresponding to the filled square in (A).

The above analysis gives little insight into the eco-evolutionary dynamics when this equilibrium is unstable. Instability occurs via destabilization of either one (or both)

of the two sets of feedbacks (i.e., the two conditions *s*(*J*_*N+PD*_) *<* 0 and *s*(*J*_*pqN−*_) *<* 0). By examining the bifurcation patterns associated with *J*_*N+PD*_ and *J*_*pqN−*_, we determine the eco-evolutionary dynamics close to the bifurcation points at which destabilization is driven by either the feedbacks in *N*_+_, *P*, *D* or *p*, *q*, *N*_−_, respectively.

If *s*(*J*_*N+PD*_) *>* 0 and *s*(*J*_*pqN−*_) *<* 0, then feedbacks between predator density, total prey density and linkage disequilibrium drive instability (red shaded region in Figure 6). From the expression (19.7) of *J*_*N+PD*_ in Supporting Information 1, we find that the transition from stability to instability when *s*(*J*_*N*+*PD*_) = 0 always occurs via a Hopf bifurcation: the focal polymorphic equilibrium is unstable and stable cycles emerge (Supporting Information 1).

Surprisingly, but as suggested by the feedbacks driving the instability, it is the predator density, total prey density, and the linkage disequilibrium that cycle around the equilibrium (Figure 7). In these cycles, as total prey density increases, predator density follows, as in classical predator-prey dynamics, and additionally, linkage disequilibrium within the predator population also follows. By examining the phase lags in our numerical solutions, we find that the predator density is a quarter-phase lagged from the total prey density and the linkage disequilibrium is a half-phase lagged from the predator density (Figure 7). The amplitude of these cycles increases and then decreases with increasing recombination rates in an interesting pattern (Figure S2).

**Figure 7.**
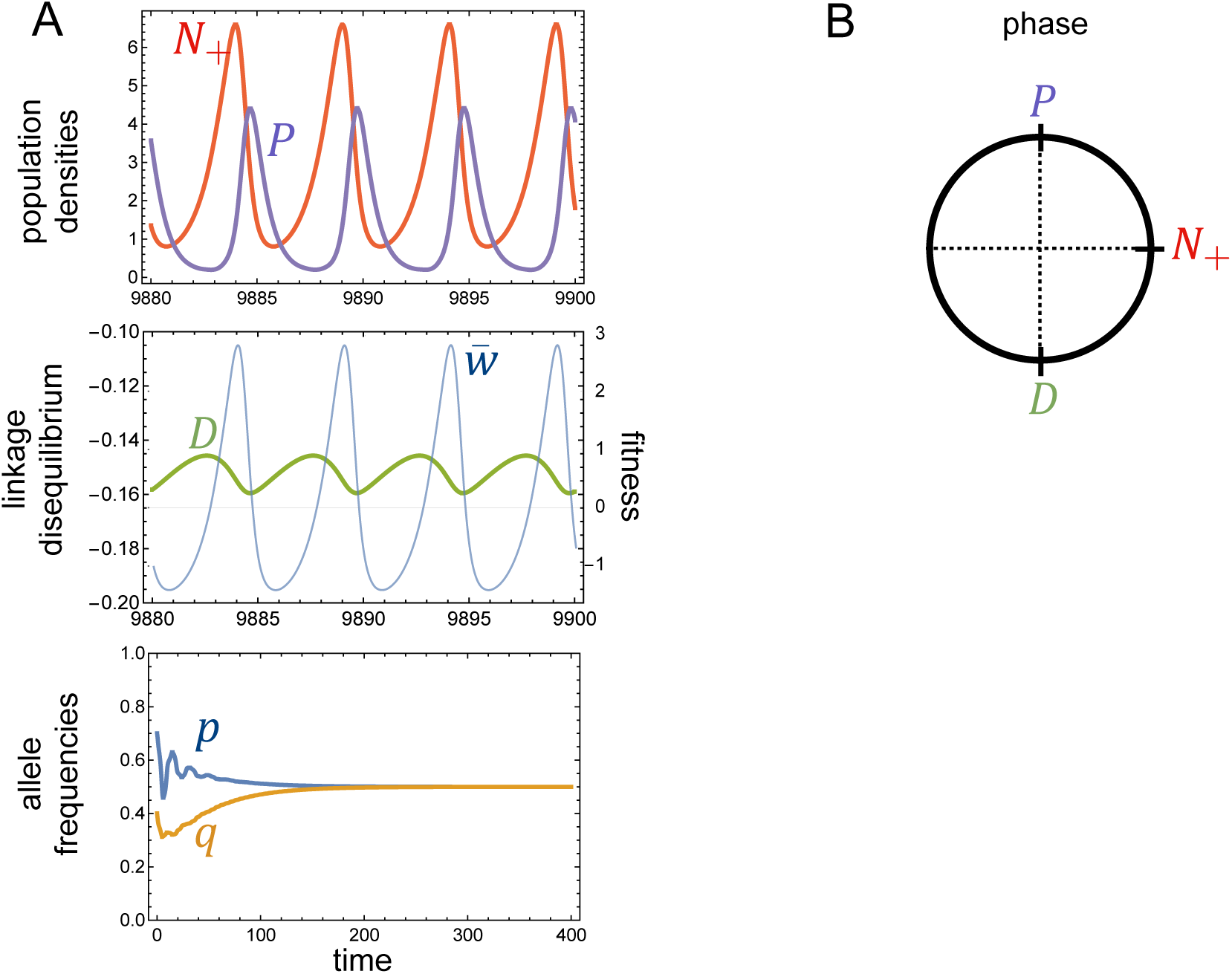
Cycling of linkage disequilibrium and predator and total prey population densities. (A) Numerical solutions corresponding to parameters of the triangle in Figure 6A (*γ*_1_ = 0.7 and *r* = 0.06). For values of *r* near the Hopf bifurcation, feedbacks cause cycles between total prey density, predator density and linkage disequilibrium (*N*_+_, *P*, and *D*), while allele frequencies approach an equilibrium value 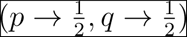 as *t* → ∞. The difference between the two prey densities converges to zero, i.e., *N*_−_ → 0 (not shown). (B) Numerics shows that the predator density and linkage disequilibrium are roughly one-quarter and three-quarters lagged, respectively, from the prey density.

Interestingly, when parameters are sufficiently near the bifurcation point, the cycles of *N*_+_, *P*, and *D* are sufficiently small that the higher-order feedbacks do not impact the stability of the equilibrium for *p*, *q*, and *N_−_*: the frequencies of the two alleles and the prey difference approach their equilibrium values. Conversely, when the cycles in *N*_+_, *P*, and *D* are large enough, these higher-order feedbacks become destabilizing and drive cycles in *p*, *q*, and *N_−_* as well (Figures 8 and S2).

**Figure 8.**
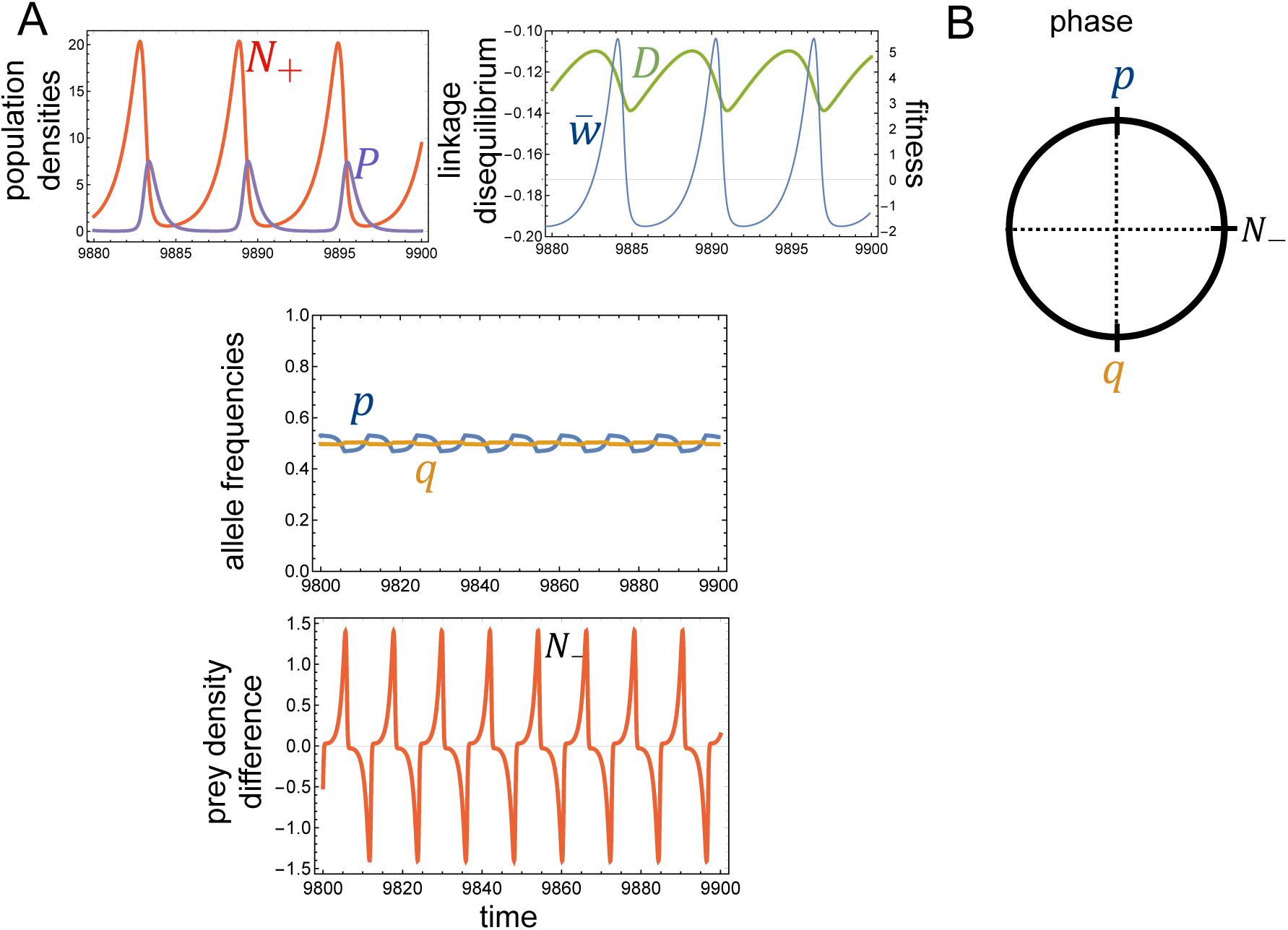
Cycling of all quantities. (A) Numerical solutions of dynamics corresponding to the star in Figure 6A (*γ*_1_ = 0.7 and *r* = 0.09). For this recombination rate, higher-order feedbacks of the cycling total prey density, predator density, and linkage disequilibrium drive cycles in the allele frequencies and difference between the two prey. (B) The two allele frequencies are out of phase, where one is quarter-lagged from the prey difference and the other is three-quarter lagged.

If *s*(*J_N_*_+*P D*_) *<* 0 and *s*(*J_pqN−_*) *>* 0, then the feedbacks between allele frequencies and the difference between the prey densities drive instability (white region below red curve in Figure 6). Numerically, we find that the transition from stability to instability of the polymorphic equilibrium point occurs via a pitchfork bifurcation when parameters are such that *s*(*J_pqN−_*) = 0. In particular, by numerically solving from various initial conditions, we observe that the fully polymorphic symmetric equilibrium becomes unstable and two new stable equilibria emerge, which are also polymorphic. Furthermore, these two new equilibria remain polymorphic only over a narrow range of parameters. Beyond this narrow range of parameters, the eco-evolutionary dynamics approach one of two alternative equilibrium points in which the locus with smaller effects fixes in one of the two alleles and all three species coexist (Figure S2). As in the case of independent loci, the initial state of the community determines the alleles that fixes and the ultimate densities of the three species.

Finally, we explore how asymmetries in the prey’s intrinsic growth rate affect doubly polymorphic equilibria. Without loss of generality, we assume that prey 1 has a higher growth rate than prey 2. We can already see from the coexistence conditions (13) that when prey 1 has a much higher growth rate, it will exclude prey 2. For small differences, we numerically solve for the nearly symmetric doubly polymorphic equilibrium 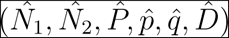 and determine the critical recombination rate below which this equilibrium is stable.

We find that as *ρ*_1_ increases, the allele frequency at the major locus, 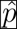, increases, and the frequency at the minor locus, 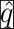, decreases (Figure S3a). Predator density increases as well (Figure S3d), and linkage disequilibrium becomes weaker, i.e., less negative (Figure S3c). For small recombination rates, the equilibrium density of prey 1 is larger than that of prey 2, as one might expect when prey 1 has larger intrinsic growth rate. Interestingly, for slightly higher recombination rates, this pattern is reversed: prey 2 is more abundant than prey 1 at equilibrium (Figure S3a). Finally, the nearly symmetric equilibrium points remain stable for higher recombination rates than when both prey are symmetric (Figure S3a).

## 5. Discussion

Eco-evolutionary feedbacks have been recognized as having important impacts on the maintenance of diversity at both the ecological and evolutionary level. For example, experimental studies with the mustard plant *Brassica nigra* showed that evolution of allelochemicals contributed to the maintenance of genetic and species diversity (Lankau and Strauss, 2007). While many of the traits in these studies are likely the outcome of complex genetic architectures, involving multiple genes and interactions between genes, most theoretical studies that account for explicit genetics do so with either a single locus (Pimental, 1968; Levin and Udovic, 1977; Wilson and Turelli, 1986; Gokhale et al., 2013; Schreiber et al., 2018), an adaptive-dynamics approach, hence assuming asexual reproduction (e.g., Abrams, 2006*a*,*b*; Rueffler et al., 2005), or a quantitative-genetics framework (Schreiber et al., 2011; Vasseur et al., 2011; Patel and Schreiber, 2015).

With only a few exceptions, the question of maintaining diversity at the ecological and evolutionary level were typically studied separately. Here, we develop a model which couples a two-locus population-genetic model to a Lotka-Volterra ecological model of three interacting species. We study how these feedbacks affect ecological diversity (the coexistence of species) and evolutionary diversity (the maintenance of polymorphism). We show that accounting for multilocus genetics of traits uncovers novel mechanisms for eco-evolutionary feedbacks between the genetics of the evolving species and the species that it interacts with.

Even with this fairly simple model, the nature of feedbacks between evolution and ecology are complex. Our approach to linearize around a double polymorphic equilibrium point enables us to identify the first-order feedback loops, which affect stability of this equilibrium. As our study demonstrates, this is a potentially powerful tool in identifying feedbacks. When prey are symmetric, our block diagonal Jacobian indicates that there are two decoupled groups of feedbacks that affect stability: (1) the total prey density (*N*_+_), the predator density (*P*), and linkage disequilibrium (*D*); and (2) the allele frequencies at both loci (*p, q*) and the difference between the prey densities (*N_−_*).

To understand the feedback in the first group, suppose the total prey density increases from equilibrium. Then, there is a two-fold response (Figure S4). On the one hand, the predator density increases, which then puts increased pressure on both prey to cause their densities to decrease, as in classical predator-prey cycles. On the other hand, as the total prey density increases, the relative fitness differences between individuals with intermediate genotypes (close to 0) and individuals with extreme genotypes (close to 1 or -1) increases (right upper loop in Figure S4). Hence, selection for intermediate genotypes is stronger, and this leads to negative linkage disequilibrium. As beneficial combinations of alleles become linked, the genotypic variance decreases and the overall fitness of the predator increases and this amplifies the negative impact the whole predator population has on the prey populations. Decreasing *N*_+_ has the opposite effects and is illustrated in the left lower loop of Figure S4.

In summary, fluctuations between periods of strong and weak selection, caused by changes in prey density, alter the genetics of the predator, which in turn drive changes in prey density. We refer to this as the fluctuating-strength-of-selection feedback (Figure S4). This novel feedback highlights the importance of considering genetic architecture in identifying the mechanisms of eco-evolutionary feedbacks, because it is mediated by the linkage disequilibrium induced by epistatic fitness effects of linked loci and frequency-dependent selection acting on them.

The mechanism behind the feedback between the allele frequencies and the difference in prey densities is as follows: suppose we perturb the system at equilibrium by increasing the density of prey 2, hence of *N*_−_ = *N*_2_ − *N*_1_ (Figure S5). Consequently, the fitness curve changes: predator individuals with higher genotypic values are more fit and there is selection for increased genotypic values. This drives the allele frequencies *p* and *q* at both loci to decrease. As they decrease, the predator becomes better at consuming prey 1 and worse at prey 2, which reduces the difference *N*_−_ in the prey densities. As *N*_−_ becomes negative, selection for low genotypic values is generated. We refer to this feedback as the alternating-direction-of-selection feedback (Figure S5).

Our model predicts that both types of eco-evolutionary feedbacks affect the maintenance of diversity by (1) promoting coexistence of species, (2) promoting the maintenance of genetic polymorphisms, and (3) driving novel eco-evolutionary cycles. In particular, here, we highlight that the alternating-direction-of-selection feedback promotes coexistence: the predator evolves to genotypes better suited for consuming the more abundant species, and this alleviates competitive pressures on the rarer prey species. Previous theoretical and experimental work has recognized this mechanism for promoting coexistence in other types of ecological interactions, including species competing via a shared predator, as in apparent competition (Schreiber et al., 2011, 2018), via direct competition (Lankau and Strauss, 2007; Vasseur et al., 2011), and intraguild predation (Ellner and Becks, 2011; Patel and Schreiber, 2015).

While eco-evolutionary feedbacks have been recognized to promote the coexistence of species, there has been significantly less attention to how they influence the maintenance of intraspecific genetic variation. Consistent with previous work on polymorphisms in two-locus models with stabilizing selection, we show that if selection is weak relative to recombination, the double polymorphism, which always has negative linkage disequilibrium, is unstable (Hastings and Hom, 1990). If selection is strong, it is stable (Gavrilets and Hastings, 1993; Bürger, 2000). Whereas selection for intermediate genotypes when both prey are present facilitates the maintenance of polymorphism, recombination breaks down the linkage of beneficial alleles. The eco-evolutionary feedbacks in our model have no qualitative effects on these results in the limits of strong or weak selection or, equivalently, in the limits of weak or strong recombination.

For intermediate recombination rates, we have identified that the two feedbacks affect the stability of the doubly polymorphic equilibrium in our model in different ways, which we infer from their effects on the Jacobian. In addition to promoting the coexistence of the two prey (discussed in a previous paragraph), the alternating-direction-of-selection feedback supports stability of the symmetric equilibrium, hence of a full polymorphism, for higher recombination rates than for the purely evolutionary dynamics with fixed population densities (compare the blue curve with the grey dashed curve in Figure 5a).

This alternating-direction-of-selection feedback resembles a previously recognized mechanism in one-locus and multilocus models that enables the maintenance of polymorphisms: certain forms of externally-driven temporal fluctuations in the environment (for example, through seasons), in which the optimal genotype changes in time, have been shown to maintain genetic variation, especially in concert with epistasis or dominance (e.g., Haldane and Jayakar, 1963; Ellner and Sasaki, 1996; Bürger and Gimelfarb, 2002; Wittmann et al., 2017).

In contrast, the fluctuating-strength-of-selection-feedback generates a novel type of eco-evolutionary cycling. Previous work shows that when considered separately from one another, the ecological equations of our model do not exhibit cycling (Takeuchi and Adachi, 1983) and neither do the evolutionary equations with either constant fitnesses (Gavrilets and Hastings, 1993; Bürger and Gimelfarb, 1999) or with negative frequencydependent selection (Bürger and Gimelfarb, 2004). Interestingly, in one class of cyclic dynamics we found, which emerges directly from the Hopf bifurcation (Figure 7), the difference between the two prey and the allele frequencies are fixed and only the total prey density, predator density and linkage disequilibrium cycle. Empirically, if we observe only allele frequencies of one or both loci independently at the population level, we would fail to uncover any eco-evolutionary feedback. If these cycles are large enough, then they give rise to cycles in the difference between the prey densities and the allele frequencies, generating a second class of cycles (Figure 8).

Two previous studies examined the eco-evolutionary dynamics in apparent competition, in which the predator was evolving in a single locus. Our results are consistent with results from Schreiber et al. (2018) when one locus has no effect on the genotype (*γ*_1_ = 0 or *γ*_2_ = 0). If, additionally, the intrinsic growth of both prey is sufficiently faster than the predator’s consumption, then we recover the model considered in Wilson and Turelli (1986), who pointed out that feedbacks enable the maintenance of a single-locus polymorphism, via a similar mechanism as our alternating-directions-of-selection feedback. There are many real biological examples in which predator and prey have similar generation times, implying that their respective biological rates are on similar time scales. Furthermore, many ecologically relevant traits are complex and the outcome of multiple loci with different effects. Accounting for these additional biological complexities may uncover new mechanisms of eco-evolutionary feedbacks and cyclic dynamics not previously recognized.

Previous work has highlighted that the shape of evolutionary trade-offs have important implications for eco-evolutionary dynamics (Rueffler et al., 2005; Schreiber et al., 2011; Patel and Schreiber, 2015). Indeed, the trade-offs influence the two feedbacks we identify here. With less concave trade-offs (smaller *s* and larger *θ*), changes in the allele frequencies result in a relatively larger change in the difference between the two prey densities. This has a destabilizing effect on the equilibrium via the alternating-direction-of-selection feedback. In contrast, less concave trade-offs have a stabilizing effect via the fluctuating-strength-of selection feedback. By lowering the fitness of the predator, there is a lower potential for relative fitness differences, which leads to only weak selection for intermediate genotypes and diminishes the feedback on linkage disequilibrium. Altogether, for less concave trade-offs, feedbacks between the allele frequencies and the prey difference become more destabilizing while feedbacks between the predator density, total prey density and linkage disequilibrium become more stabilizing. Hence, there is an optimal trade-off for stability (Figure S6).

In this work, we study the simplest possible multilocus model, with two diallelic loci, to account for more complex genetic architecture than in previous studies. We find novel eco-evolutionary feedbacks between genetics of the evolving species and the community that previously were not recognized but are important for the stability of polymorphisms. An important avenue for future work is to understand how eco-evolutionary dynamics with more than two loci influence the coexistence and the maintenance of polymorphisms. Do we recover similar types of feedbacks when we consider more than two loci? Are there additional novel types of eco-evolutionary feedbacks? Due to the complexity of multilocus dynamics, we suggest that gaining insight into these questions will require computational techniques.

## Acknowledgements

This work was supported by the Austrian Science Fund (FWF) through Grant P25188-N25.

## Appendix 1: Full differential equation model

In this appendix, we give the full expressions of the differential equation model, including explicit expressions for all partial derivatives in equation (7). We define

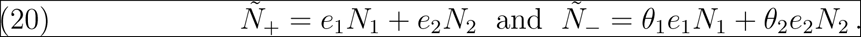

From (8) and (9), we deduce easily, by substituting (2), (1), (10), and (11),

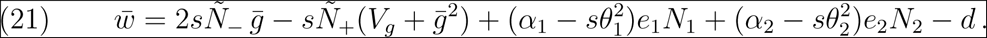

Therefore, we obtain

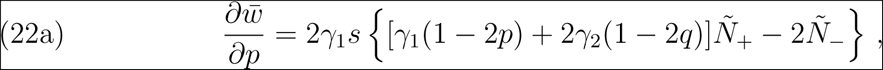

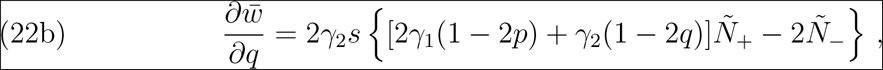

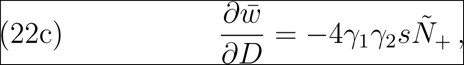

and, by substituting this into (7),

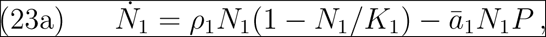

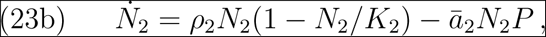

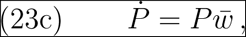

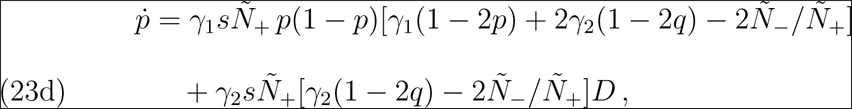

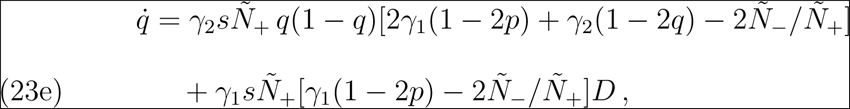

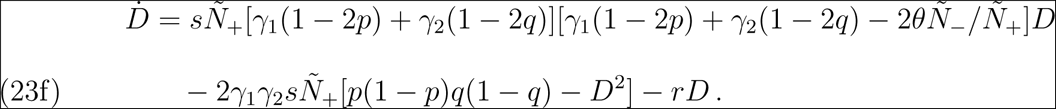

The equations for 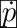 and 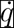 show that *sÑ*_+_ is a measure of the strength of density-dependent selection. In addition, the first term in the second line for 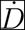 is always negative due to (6b). This reflects the well known fact that stabilizing selection induces negative linkage disequilibrium through its epistatic effects.

## Appendix 2: Coexistence of two prey

In this appendix, we derive the conditions for persistence of the two prey species. Heuristically, the two prey species will persist if each can invade, i.e., has, on average, positive per-capita growth rate, in a community when it is absent.

If prey 1 is absent, i.e., *N*_1_ = 0, then solutions to our differential equation model converge to the equilibrium point

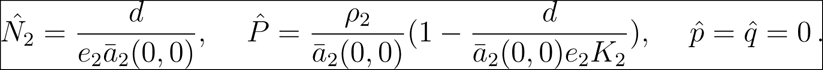

Biologically, the predator evolves to optimize on prey 2 and the predator and prey coexist at the corresponding equilibrium. Then, the per-capita growth rate of prey 1 at this equilibrium is

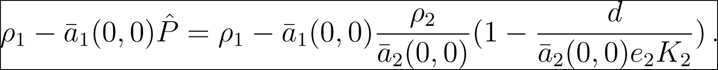

The condition for this to be positive, i.e., for prey 1 to be able to invade, simplifies to (13a). Analogously, the condition for prey 2 to be able to invade when initially absent is (13b).

### Supporting Information 1

In this supporting information, we provide details on the stability analysis of the fully polymorphic equilibrium when the prey are symmetric, i.e., we assume condition (16). We give rigorous results and also point out which results come from numerical analysis.

With the substitution (17), so that *Ñ*_+_ = *eN*_+_ and *Ñ*_+_ = *θeN_−_*, the dynamics (23) becomes

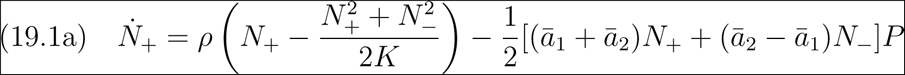

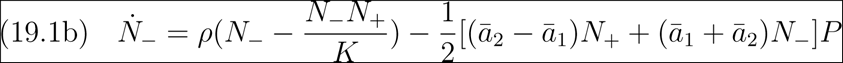

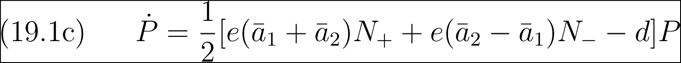

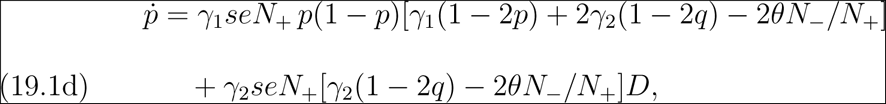

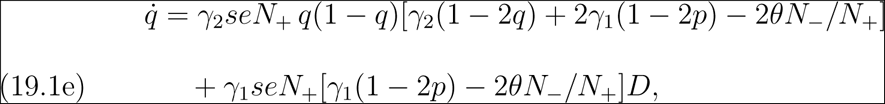

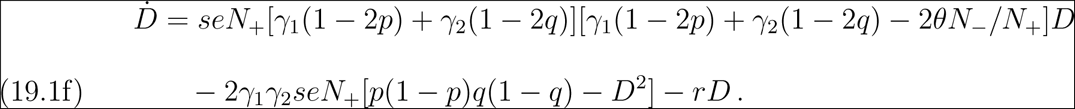

A straightforward calculation based on (5) shows that for given and fixed *N*_+_ and *N_−_*, the evolutionary equations (19.1d), (19.1e), and (19.1f), become equivalent to the equations for the well known quadratic-stabilizing-selection model with an arbitrary optimum and frequency- and density-independent selection (Gavrilets and Hastings 1993; Bürger 2000, pp. 213–216). For independent loci (*D* = 0), the model of Hastings and Hom (1990) is obtained; its equilibrium and stability structure was derived analytically. Although a fully polymorphic equilibrium may exist, it is never stable. For loci recombining at rate *r* and if *θN_−_* = 0 in (19.1d), (19.1e), and (19.1f), the quadratic-stabilizing-selection model with a symmetric optimum is obtained. In this model, for given *s >* 0 and *γ*_1_ ≠ *γ*_2_, there exist critical recombination rates 0 *< r*_1_ *< r*_2_, such that the symmetric two-locus polymorphism is globally asymptotically stable if *r < r*_1_, two asymmetric two-locus polymorphisms exist and are stable if *r*_1_ *< r < r*_2_, and one or both loci become fixed if *r > r*_2_ (Bürger and Gimelfarb 1999; Bürger 2000, pp. 204–208).

Back to the full dynamics (19.1), if 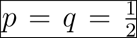, *N*_−_ = 0, *P* > 0, and the prey are symmetric, we observe that 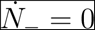 implies 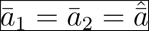 and obtain

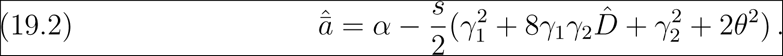

Hence, 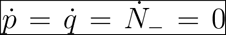 in (19.1). Using these values for *p*, *q*, and *N*_−_, we find the linkage disequilibrium (18b) at the symmetric equilibrium by solving 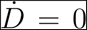. In the case of independent loci, we automatically have *D*^ˆ^ = 0, as expected by taking the limit *r → ∞* in (18b). In each case, explicit expressions for the equilibrium values 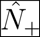 and 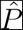 are obtained as follows. By solving 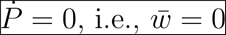, we find

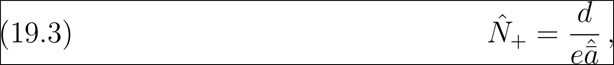

and from 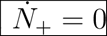, we obtain

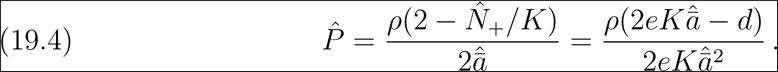

If the loci are independent, then 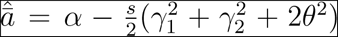, and substitution into (19.3) and (19.4) yields very simple expressions.

With linked loci (*r ≥* 0), the situation is more complicated because then 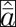 depends on 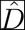, given by (18b), which in turn depends on 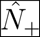. Therefore, a quadratic equation for 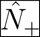 is obtained. In Supporting Information 2, we show that the unique admissible solution is given

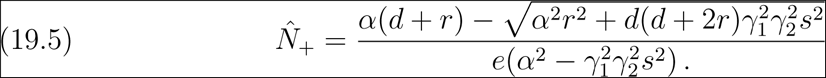

The explicit expression for 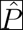 is much more complicated and given in Supporting Information 2.

We can show that 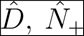, and 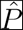 are increasing functions in *r*, and 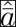 is decreasing (but recall that 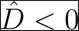). However, because 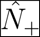 (for independent loci) and *d/*(*ea*_1_ + *esγ*_1_*γ*_2_) (for completely linked loci), where 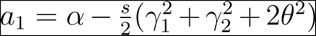, the dependence on *r* is very weak if *α* is large (see also Figures S2 and S3).

For an equilibrium point, we define the stability modulus as the largest real part of the set of eigenvalues of the Jacobian, *J*, and label it *s*(*J*). Stability of an equilibrium point requires that *s*(*J*) *<* 0. The Jacobian of the unique equilibrium for (19.1) in the form 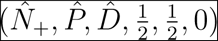 takes the block diagonal form

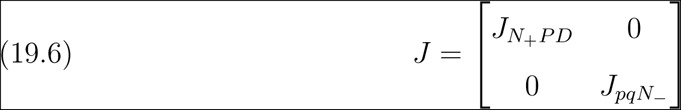

with

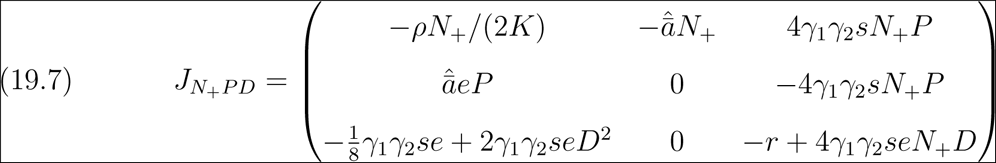

and

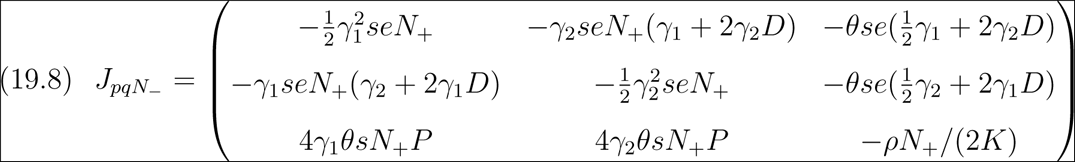

This block diagonal structure makes it easier to evaluate stability, since *s*(*J*) = max*{s*(*J_pqN−_*)*, s*(*J_N_*_+*P D*_)*}*. Hence, the equilibrium point is stable if and only if *s*(*J_pqN−_*) *<* 0 and *s*(*J_N_*_+*P D*_) *<* 0.

Generally, for a 3 *×* 3 Jacobian, *J*, it is straightforward to use the Routh-Hurwitz criteria to determine if *s*(*J*) *<* 0, i.e., the corresponding equilibrium is stable. Let 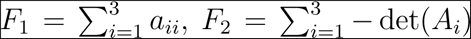, and *F*_3_ = det(*J*), where *a*_*ii*_ are diagonal terms and *A_i_* is the submatrix of *J* with the *i^th^* row and column deleted. In words, these are the trace, sum of the principal minors, and the determinant of *J*. Then, the number of eigenvalues with positive real part are the number of sign changes in the sequence *{*1*, −F* 1*, −F* 1(*F* 1*F* 2 + *F* 3)*, −F* 3*}* (Gantmacher 1998). In particular, *s*(*J*) *<* 0 if and only if *F*_1_ *<* 0*, F*_3_ *<* 0 and *F*_1_*F*_2_ + *F*_3_ *>* 0. Using this criterion, we are able to determine the stability of the symmetric equilibrium (18a) with *D*^ˆ^ = 0 as well as with (18b) for a number of special cases.

For *J_N_*_+*P D*_, we get

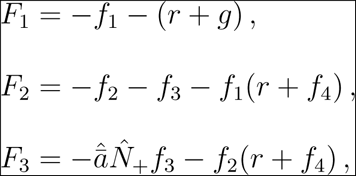

where

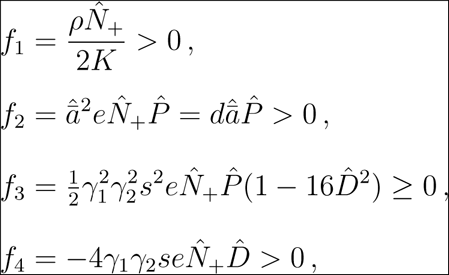

and we have used 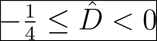. It follows immediately that *F*_1_ < 0, *F*_2_ <0, and *F*_3_ *<* 0.

Next, we examine *F*_1_*F*_2_ + *F*_3_ *>* 0. A simple calculation yields

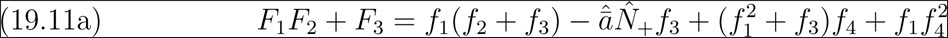

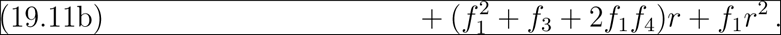

If *r* = 0, then 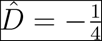 and *f*_3_ = 0. It follows that 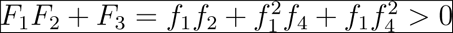. By continuity, *F*_1_*F*_2_ + *F*_3_ is positive for sufficiently small *r*, whence *s*(*J_N_*_+*P D*_) *<* 0 holds for sufficiently small *r*. If 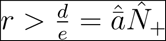, then *F*_1_*F*_2_ +*F*_3_ *>* 0 because 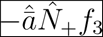 is the only negative term in (19.11), and the right-hand side of (19.11a) is positive if *r >* 0; in fact, it tends to infinity if *r → ∞* (because 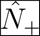 and 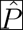 remain bounded), and so *s*(*J_N_*_+*P D*_) *<* 0. By plotting, we observe that *F*_1_*F*_2_ + *F*_3_ is often negative for intermediate values of *r*(Supporting Information 2).

The corresponding expressions for *J_pqN−_*are

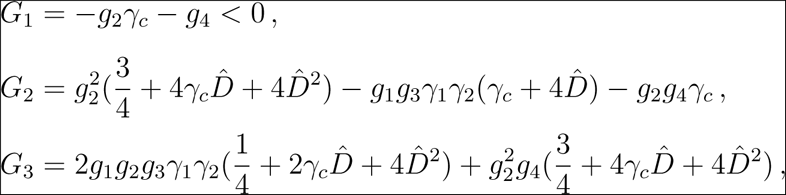

where

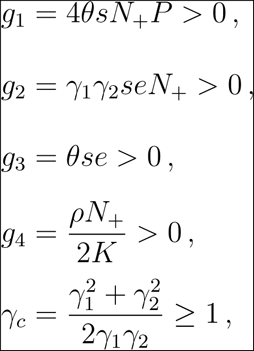

Obviously, we have *G*_1_ *<* 0. Hence, the remaining stability conditions to ensure that *s*(*J_pqN−_*) *<* 0 are *G*_3_ *<* 0 and *G*_1_*G*_2_ + *G*_3_ *>* 0.

Altogether, we have *s*(*J*) *<* 0 if and only if *F*_1_*F*_2_+*F*_3_ *>* 0, *G*_3_ *<* 0, and *G*_1_*G*_2_+*G*_3_ *>* 0. We use these conditions to determine the stability for the symmetric equilibrium under various assumptions.

**If loci are independent or sufficiently loosely linked, the symmetric equilibrium is unstable.**

First, assume 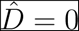. Then we obtain

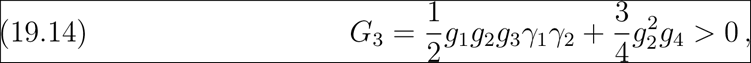

which yields *s*(*J_pqN−_*) *>* 0, hence instability. Because the symmetric equilibrium (18a) with (18b) converges to the symmetric equilibrium with 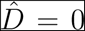 as *r → ∞*, it is also unstable for sufficiently large *r*.

**If** *γ*_1_ ≠ *γ*_2_**, the symmetric equilibrium** (18) is unstable.

If *γ*_1_ = *γ*_2_, then *γ_c_* = 1 and, by examining the two quadratic functions of 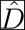 in the expression of *G*_3_, we observe that *G*_3_ *≥* 0 for all 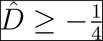(equality holds only if 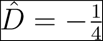), hence instability.

**If** *γ*_1_ ≠ *γ*_2_**, the symmetric equilibrium** (18) **is asymptotically stable for sufficiently small** *r***, and it is unstable for sufficiently large** *r*.

If *r* = 0, then 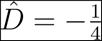 and

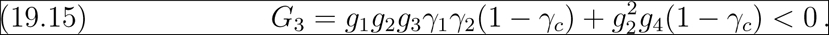

Additionally, *F*_1_*F*_2_ + *F*_3_ *>* 0 (see above) and *G*_1_*G*_2_ + *G*_3_ *>* 0 (Supporting Information 2). Therefore, *s*(*J*) *<* 0 if *r* = 0. By continuity, *s*(*J*) *<* 0 holds for sufficiently small *r*, whence the symmetric equilibrium is stable. As *r → ∞*, 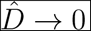. Therefore, by (19.14), *G*_3_ becomes positive for sufficiently large *r*, which establishes the assertion.

These findings parallel and extend those outlined above for the asymptotic stability of the symmetric equilibrium in the absence of frequency- and density-dependent selection.

---

**Figure S1.**
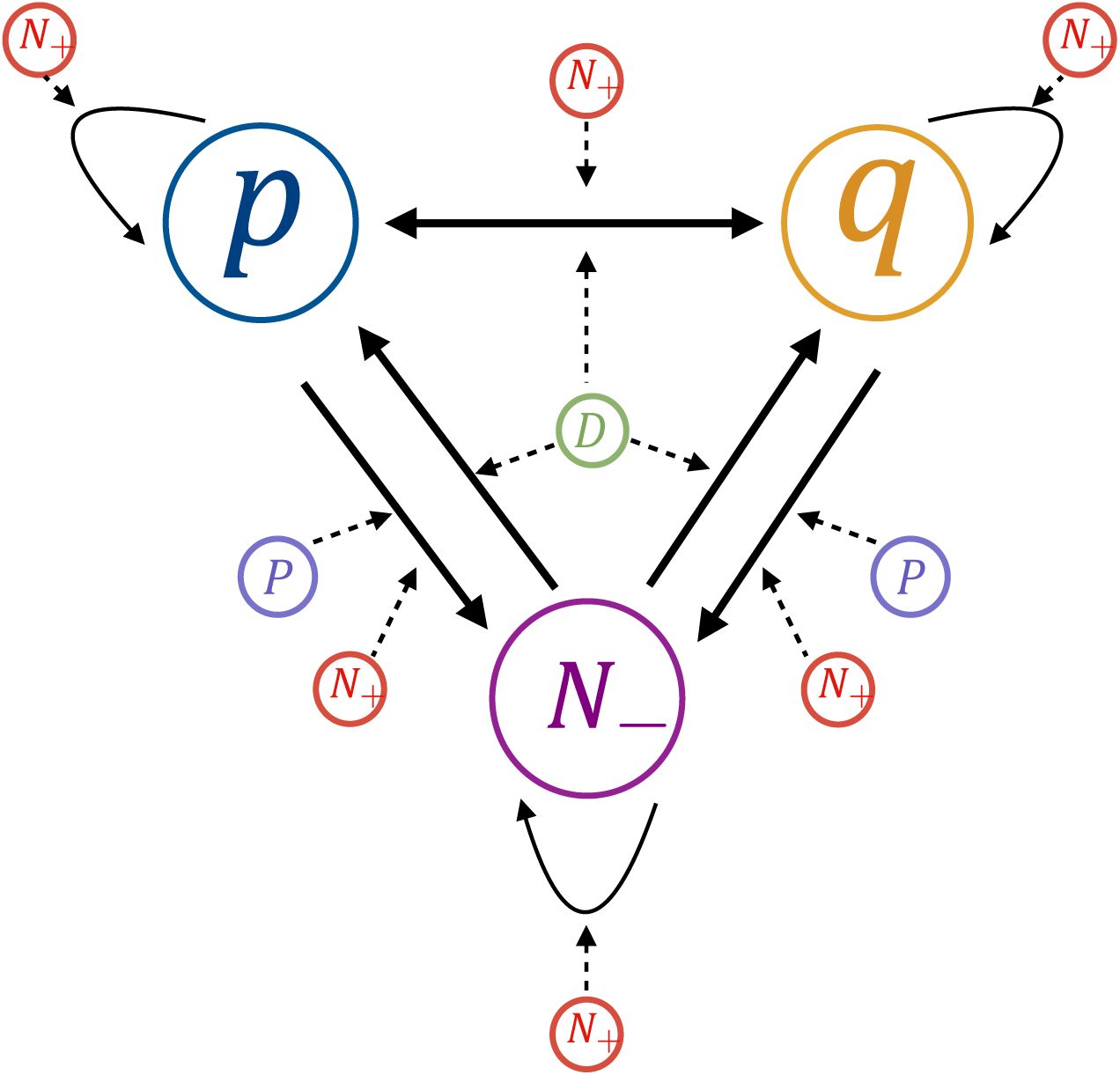
Higher-order feedbacks impacting stability of the asymmetric double polymorphic equilibrium. The first-order feedbacks (solid arrows) between prey density difference and both allele frequencies are determined from non-zero entries of the Jacobian matrix (19.8). For example, if *J_ij_ >* 0, then there is a positive influence of *j* on *i*. The higher-order feedbacks (dotted arrows) are the effects of total prey density, predator density, and linkage disequilibrium on entries of the Jacobian. For example, if *J_ij_* is a function of variable *k*, then the variable *k* affects how strongly *j* influences *i*. There are no higher-order feedbacks of prey density difference and both allele frequencies on total prey density, predator density, and linkage disequilibrium.

**Figure S2.**
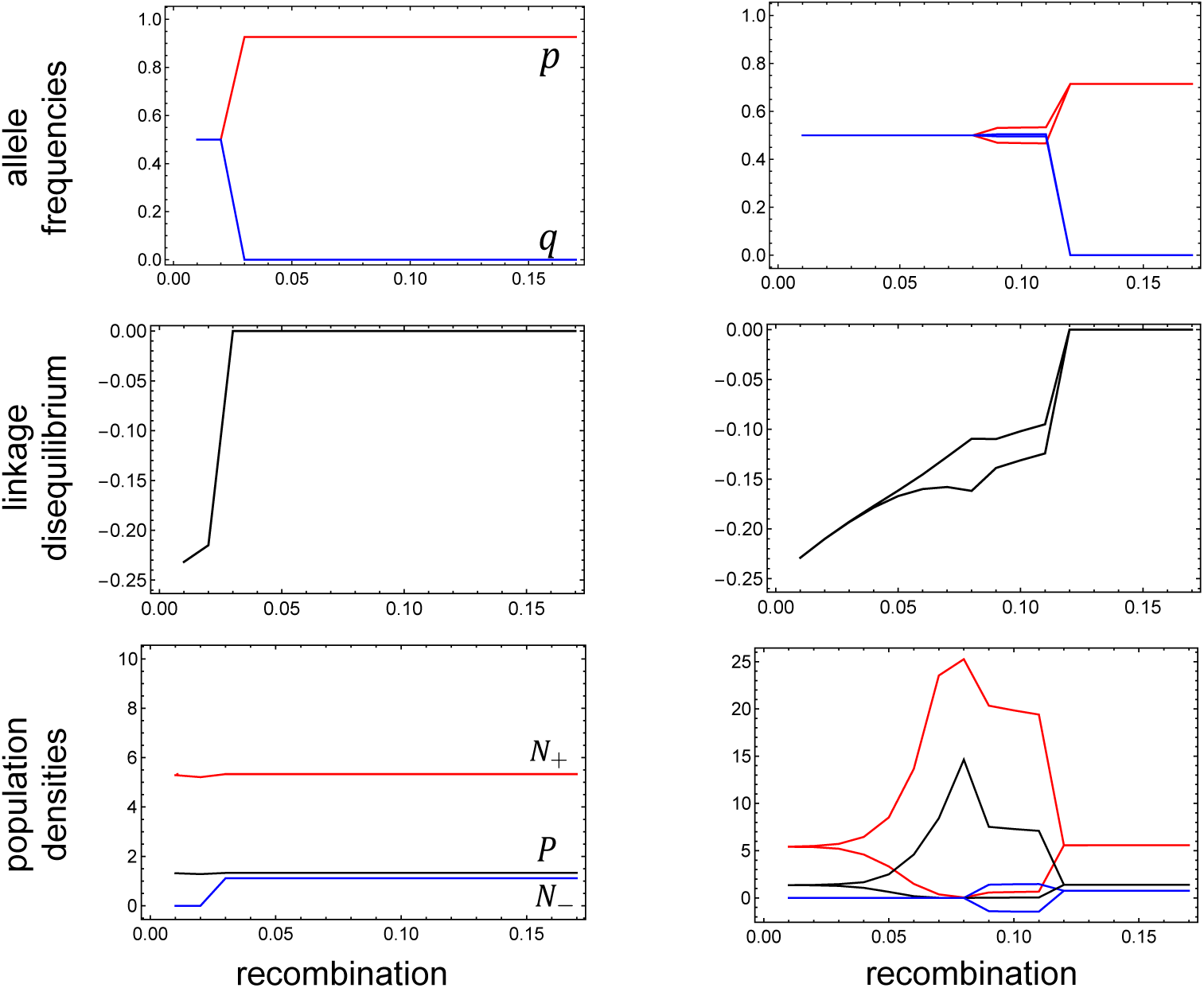
Bifurcations from the symmetric equilibrium (18) as a function of the recombination rate. The left and right panels are for parameters (see below) corresponding to the left and right vertical lines in Figure 6A, respectively. In the left panel, the symmetric equilibrium becomes unstable by a pitchfork bifurcation and a pair of asymmetric, stable equilibria emerges. They move quickly to the boundary. For the equilibrium shown here, *q* → 0 and *p* to a value close to 0.9. For the other equilibrium (not shown), *q* → 1 and *p* to approximately 0.75. In the right panel, curves split because stable cycling occur. The upper (lower) curve shows the maximum (minimum) of the respective quantity. The first bifurcation (at *r* ≈ 0.05) involves only *N*_+_, *P*, and *D* (Note: there seems to be cycling before this bifurcation due to really long transients). At *r ≈* 0.085, a second bifurcation occurs by which the allele frequencies and *N_−_* start cycling, too. Between *r* ≈ 0.11 and *r* ≈ 0.12 apparently two cycles in the allele frequencies exist (of which only one is shown in the right upper panel), which quickly move to the boundary. For *r* ⪆ 0.12, a pair of one-locus polymorphisms is stable. Parameters in common are *α*_1_ = *α*_2_ = 1, −*θ*_1_ = *θ*_2_ = 1.1, *s* = 0.2, *e*_1_ = *e*_2_ = 0.5, *K*_1_ = *K*_2_ = 1500, *ρ*_1_ = *ρ*_2_ = 1, *d* = 2. In the left panel, we have *γ*_1_ = 0.54, and the right panel, *γ*_1_ = 0.7. The minimum and maximum were recorded in between time step 9000 and 9900.

**Figure S3.**
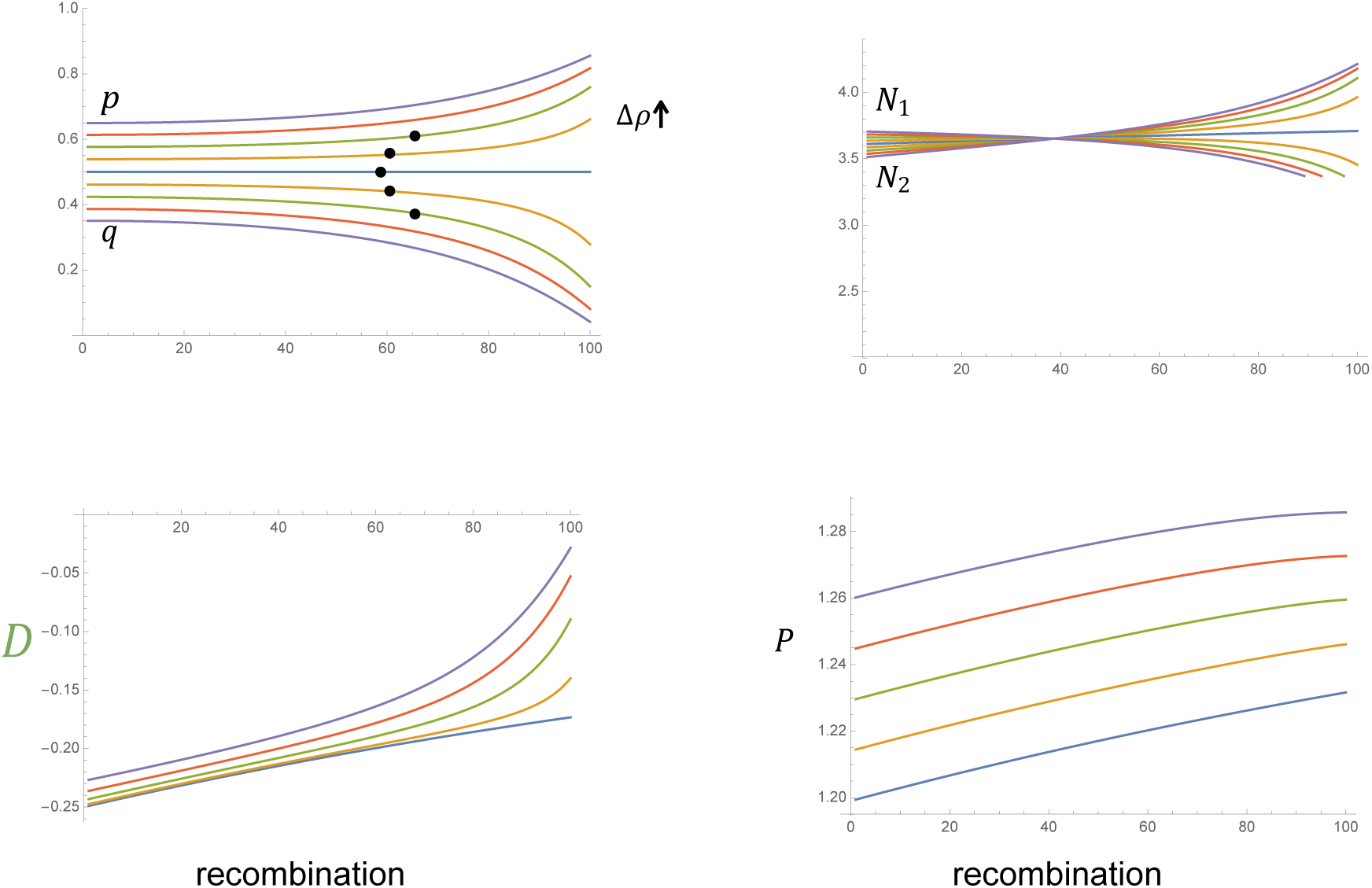
Bifurcation of equilibria for asymmetric prey. The equilibrium values for *p, q, N*_1_*, N*_2_*, D,* and *P* are shown for increasing recombination rates and varying asymmetries of the prey, measured by the difference between the intrinsic growth rates of the two prey, i.e., ∆*ρ* = *ρ*_1_ − *ρ*_2_. Parameters are *e* = 0.5*, d* = 3*, K* = 1000*, s* = 0.3*, α* = 1.2*, θ* = 1.1*, γ*_1_ = 0.6 and *ρ*_2_ = 1. The different colors are solutions for different values of *ρ*_1_, which increases from 1 to 1.1 in increments of 0.25 in the order blue, orange, green, red, purple. Dots in the first plot indicate the critical recombination rate for which stability of the equilibrium point changes from stable to unstable.

**Figure S4.**
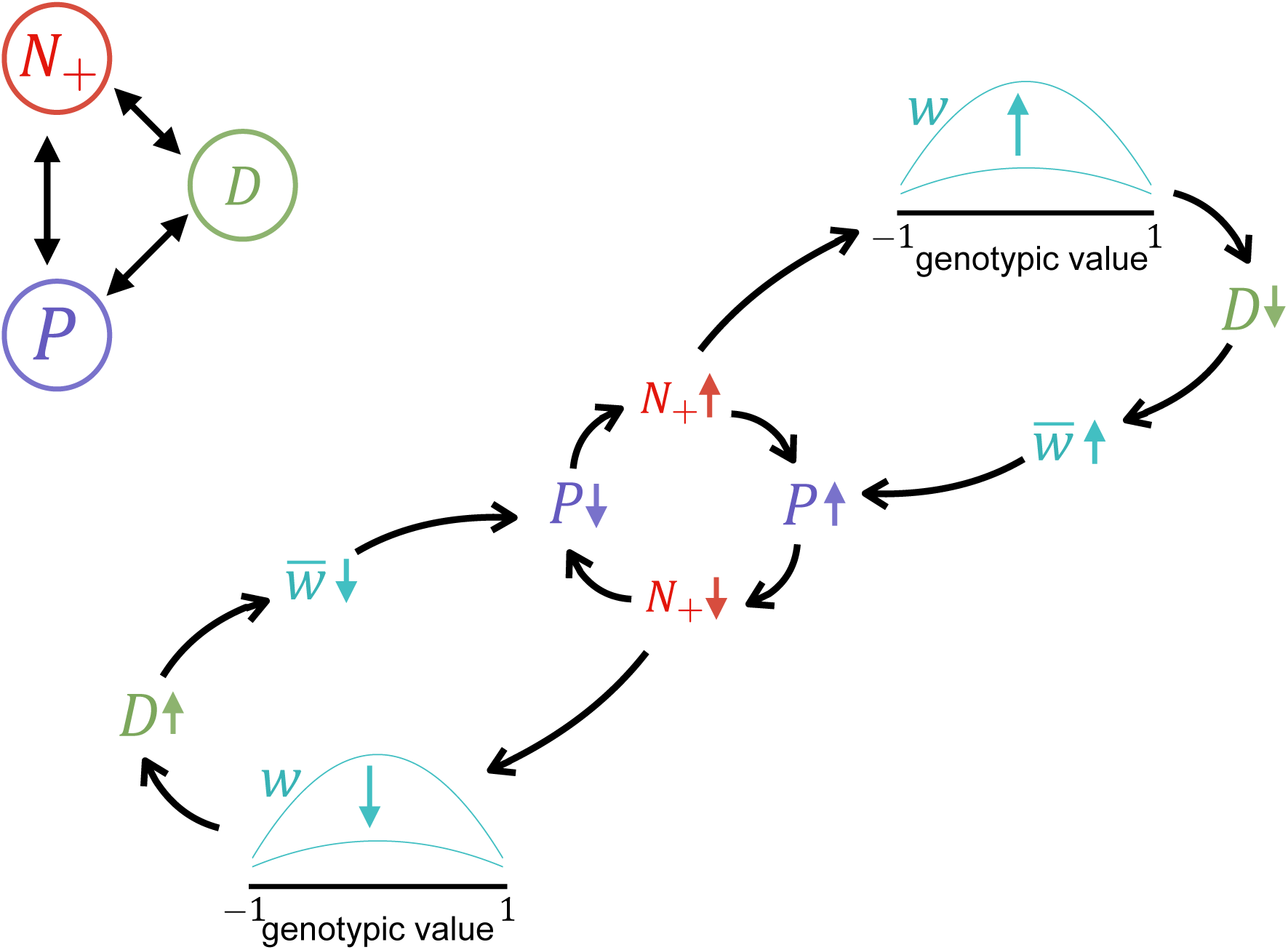
Schematic of fluctuating-strength-of-selection feedback. Increasing total prey density drives the individual predator fitness curve to be more concave, generating stronger selection for intermediate genotypes. This creates more negative linkage disequilibrium, opposing recombination, which increases the overall mean fitness of the predator population and the predator population density. Then, the total prey density decreases and this has the opposite effect on the fitness curves and linkage disequilibrium, repeating the cycle.

**Figure S5.**
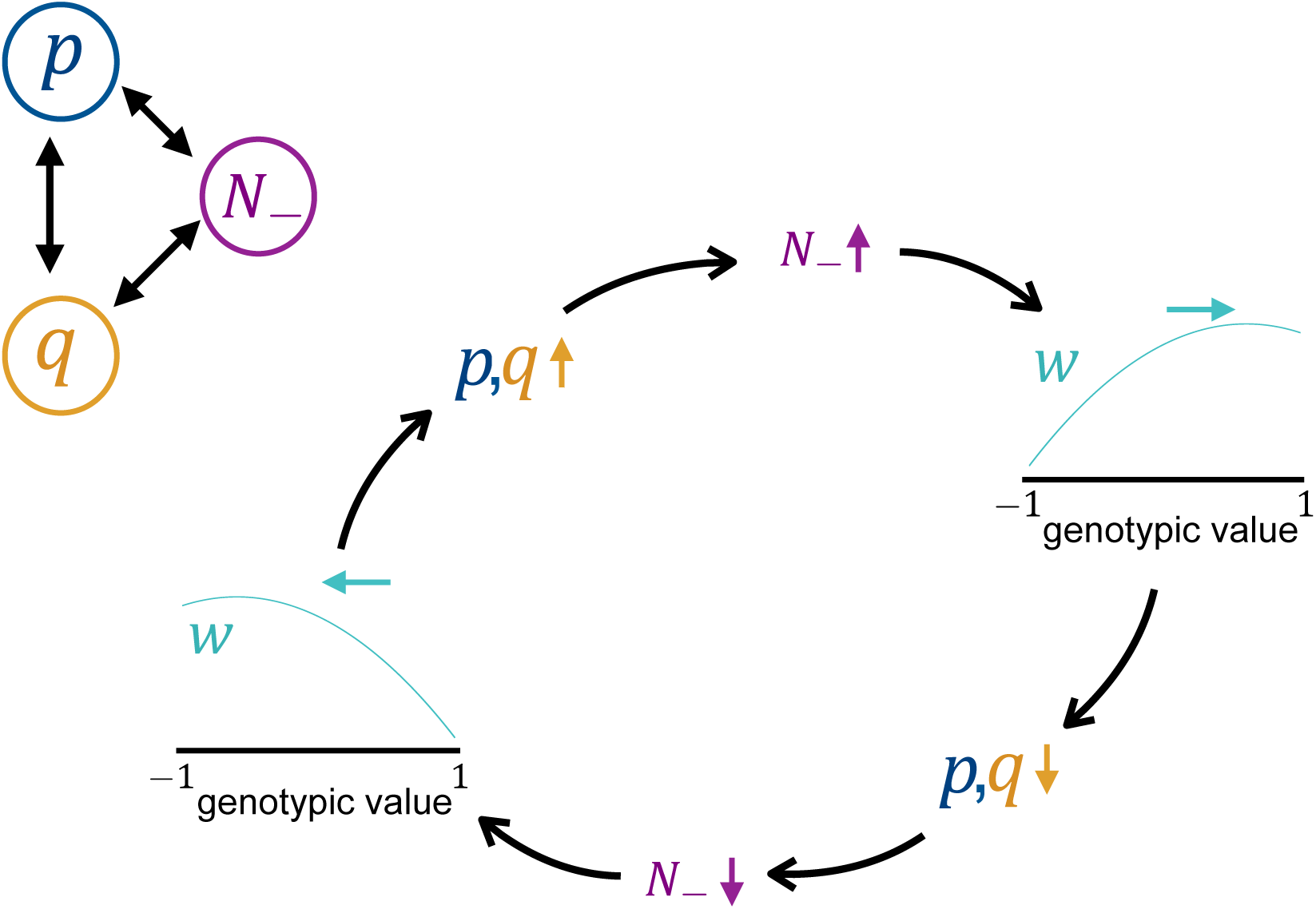
Schematic of alternating-direction-of-selection feedback. As *p* and *q* increase, the predator becomes better at attacking prey 1, so *N*_1_ decreases and *N_−_* increases. The peak of the individual predator fitness curve shifts towards higher genotypic values, which in turn drives the predator to evolve to decrease *p* and *q*. This drives *N_−_* to decrease, which shifts the individual predator fitness curve towards lower genotypic values, which drives the predator to evolve to increase *p* and *q*, repeating the cycle.

**Figure S6.**
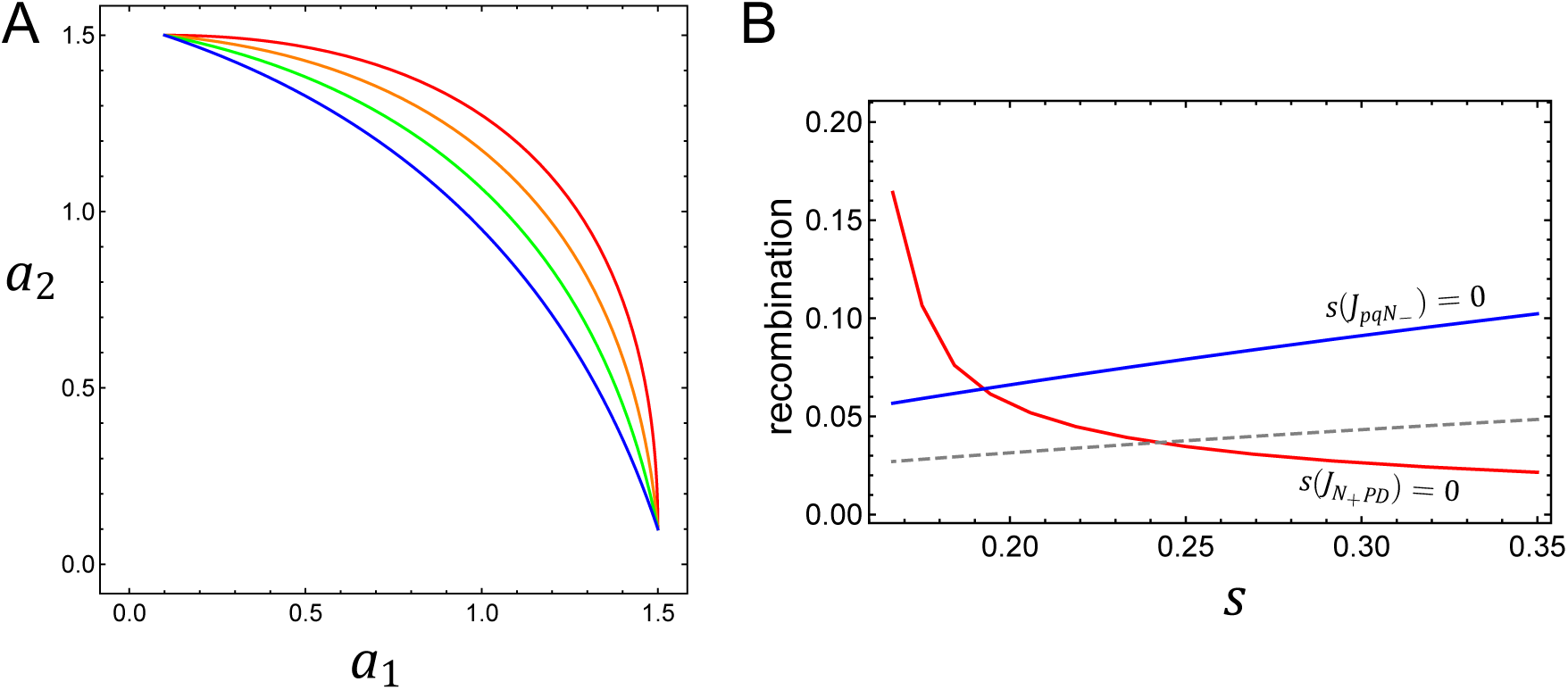
Effects of shape of trade-off on stability. (A) The attack rate trade-off curves for *s* = 0.35, 0.29, 0.23, 0.175, and *θ* = 1, 1.2, 1.5, 2 in red, orange, green, and blue, respectively. Note that *θ* was simultaneously changed with *s* in order to keep the minimum and maximum attack rate at 0.1 and 1.5, respectively. (B) The effects of this trade-off curve on the critical recombination rate for loss of stability via the two different sets of feedbacks. Parameters are *e* = 0.5*, d* = 2*, K* = 1500*, γ*_1_ = 0.66 and *ρ* = 1.

## References

Abrams, P. 2006a. The effects of switching behavior on the evolutionary diversification of generalist consumers. The American Naturalist 168:645–659.

Abrams, P. A. 2006b. Adaptive change in the resource-exploitation traits of a generalist consumer: The evolution and coexistence of generalists and specialists. Evolution 60:427–439.

Abrams, P. A. and Kawecki, T. J. 1999. Adaptive host preference and the dynamics of host-parasitoid interactions. Theoretical population biology 56:307–324.

Bürger, R., 2000. The mathematical theory of selection, recombination, and mutation. John Wiley and Sons.

Bürger, R. and Gimelfarb, A. 1999. Genetic variation maintained in multilocus models of additive quantitative traits under stabilizing selection. Genetics 152:807–820.

Bürger, R. and Gimelfarb, A. 2002. Fluctuating environments and the role of mutation in maintaining quantitative genetic variation. Genetics Research 80:31–46.

Bürger, R. and Gimelfarb, A. 2004. The effects of intraspecific competition and stabilizing selection on a polygenic trait. Genetics 167:1425–1443.

Charlesworth, B. and Charlesworth, D., 2010. Elements of evolutionary genetics. Roberts & Company.

Cortez, M. H. and Patel, S. 2017. The effects of predator evolution and genetic variation on predatorprey population-level dynamics. Bulletin of Mathematical Biology 79:1510–1538.

Ellner, S. and Sasaki, A. 1996. Patterns of genetic polymorphism maintained by fluctuating selection with overlapping generations. Theoretical Population Biology 50:31–65.

Ellner, S. P. and Becks, L. 2011. Rapid prey evolution and the dynamics of two-predator food webs. Theoretical Ecology 4:133–152.

Falconer, D. S. and Mackay, T. F., 1996. Introduction to Quantitative Genetics. Pear-son.

Gavrilets, S. and Hastings, A. 1993. Maintenance of genetic variability under strong stabilizing selection: a two-locus model. Genetics 134:377–386.

Gimelfarb, A. 1989. Genotypic variation for a quantitative character maintained under stabilizing selection without mutations: epistasis. Genetics 123:217–227.

Gokhale, C. S., Papkou, A., Traulsen, A., and Schulenburg, H. 2013. Lotka-volterra dynamics kills the red queen: population size fluctuations and associated stochasticity dramatically change host-parasite coevolution. BMC evolutionary biology 13:254.

Haldane, J. and Jayakar, S. 1963. Polymorphism due to selection of varying direction. Journal of Genetics 58:237–242.

Hastings, A. and Hom, C. L. 1990. Multiple equilibria and maintenance of additive genetic variance in a model of pleiotropy. Evolution 44:1153–1163.

Hermisson, J., Hansen, T. F., and Wagner, G. P. 2003. Epistasis in polygenic traits and the evolution of genetic architecture under stabilizing selection. The American Naturalist 161:708–734.

Holt, R. 1977. Predation, apparent competition, and the structure of prey communities. Theoretical Population Biology 12:197–229.

Holt, R. D. and Bonsall, M. B. 2017. Apparent competition. Annual Review of Ecology, Evolution, and Systematics 48:447–471.

Johnson, T. and Barton, N. 2005. Theoretical models of selection and mutation on quantitative traits. Philosophical Transactions of the Royal Society of London B: Biological Sciences 360:1411–1425.

Lande, R. 1976. Natural selection and random genetic drift in phenotypic evolution. Evolution 30:314–334.

Lankau, R. A. and Strauss, S. Y. 2007. Mutual feedbacks maintain both genetic and species diversity in a plant community. Science 317:1561–1563.

Levin, S. and Udovic, J. 1977. A mathematical model of coevolving populations. The American Naturalist Pages 657–675.

Menge, B. A. 1995. Indirect effects in marine rocky intertidal interaction webs: patterns and importance. Ecological monographs 65:21–74.

Morris, R. J., Lewis, O. T., and Godfray, H. C. J. 2004. Experimental evidence for apparent competition in a tropical forest food web. Nature 428:310.

Muller, C. a. and Godfray, H. 1997. Apparent competition between two aphid species. Journal of Animal Ecology Pages 57–64.

Nagylaki, T. 1989. The maintenance of genetic variability in two-locus models of stabilizing selection. Genetics 122:235–248.

Nagylaki, T. and Crow, J. F. 1974. Continuous selective models. Theoretical Population Biology 5:257–283.

Patel, S. and Schreiber, S. J. 2015. Evolutionarily driven shifts in communities with intraguild predation. The American Naturalist 186:E98-E110.

Phillips, P. C. 2008. Epistasisthe essential role of gene interactions in the structure and evolution of genetic systems. Nature Reviews Genetics 9:855.

Pimental, D. 1968. Population regulation and genetic feedback. Science Pages 1–7.

Pontz, M., Hofbauer, J., and Bürger, R. 2018. Evolutionary dynamics in the two-locus two-allele model with weak selection. Journal of mathematical biology 76:151–203.

Rueffler, C., Van Dooren, T. J., and Metz, J. A. 2005. The evolution of resource specialization through frequency-dependent and frequency-independent mechanisms. The American Naturalist 167:81–93.

Schmitt, R. J. 1987. Indirect interactions between prey: apparent competition, predator aggregation, and habitat segregation. Ecology 68:1887–1897.

Schreiber, S. J., Bürger, R., and Bolnick, D. I. 2011. The community effects of pheno-typic and genetic variation within a predator population. Ecology 92:1582–1593.

Schreiber, S. J., Patel, S., and terHorst, C. 2018. Evolution as a coexistence mechanism: Does genetic architecture matter? The American Naturalist 191:407–420.

Settle, W. and Wilson, L. 1990. Invasion by the variegated leafhopper and biotic interactions: parasitism, competition, and apparent competition. Ecology 71:1461–1470.

Takeuchi, Y. and Adachi, N. 1983. Existence and bifurcation of stable equilibrium in two-prey, one-predator communities. Bulletin of mathematical Biology 45:877–900.

Turelli, M. and Barton, N. 2006. Will population bottlenecks and multilocus epistasis increase additive genetic variance? Evolution 60:1763–1776.

Vasseur, D. A., Amarasekare, P., Rudolf, V. H., and Levine, J. M. 2011. Eco-evolutionary dynamics enable coexistence via neighbor-dependent selection. The American Naturalist 178:E96–E109.

Vasseur, D. A. and Fox, J. W. 2011. Adaptive dynamics of competition for nutritionally complementary resources: character convergence, displacement, and parallelism. The American Naturalist 178:501–514.

Wilson, D. S. and Turelli, M. 1986. Stable underdominance and the evolutionary invasion of empty niches. The American Naturalist 127:835–850.

Wittmann, M. J., Bergland, A. O., Feldman, M. W., Schmidt, P. S., and Petrov, D. A. 2017. Seasonally fluctuating selection can maintain polymorphism at many loci via segregation lift. Proceedings of the National Academy of Sciences 114:E9932–E9941.

Wright, S. 1935. Evolution in populations in approximate equilibrium. J. Genetics 30:257–266.

